# Rearing environment persistently modulates the phenotype of mice

**DOI:** 10.1101/2022.02.11.480070

**Authors:** Ivana Jaric, Bernhard Voelkl, Melanie Clerc, Marc W. Schmid, Janja Novak, Marianna Rosso, Reto Rufener, Vanessa Tabea von Kortzfleisch, S. Helene Richter, Manuela Buettner, André Bleich, Irmgard Amrein, David P. Wolfer, Chadi Touma, Shinichi Sunagawa, Hanno Würbel

## Abstract

The phenotype of an organism results from its genotype and the influence of the environment throughout development. Even when using animals of the same genotype, independent studies may test animals of different phenotypes, resulting in poor replicability due to genotype-by-environment interactions ^1–4^. Thus, genetically defined strains of mice may respond differently to experimental treatments depending on their rearing environment ^5^. However, the extent of such phenotypic plasticity and its implications for the replicability of research findings have remained unknown. Here, we examined the extent to which common environmental differences between rearing facilities modulate the phenotype of genetically homogeneous (inbred) mice. We conducted a comprehensive multi-center study, where inbred mice from the same breeding stock were reared in five different facilities throughout early life and adolescence, before being transported to a single test laboratory. We found persistent effects of rearing facility on the composition and heterogeneity of the gut microbial community. These effects were paralleled by persistent differences in body weight and in the behavioural phenotype of the mice. Furthermore, we show that common variation among rearing facilities is strong enough to influence epigenetic patterns in neurons at the level of chromatin organization. We detected changes in chromatin organization in the regulatory regions of genes involved in nucleosome assembly, neuronal differentiation, synaptic plasticity and regulation of behavior. Our findings demonstrate that common environmental differences between rearing facilities may produce facility-specific phenotypes, from the molecular to the behavioural level. We expect our findings to stimulate further research into the mechanisms and drivers of these epigenetic changes mediated by the laboratory environment. Furthermore, they highlight an important limitation of inferences from single-laboratory studies and a need to account for the animals’ environmental background in study design to produce robust and replicable findings.

## Introduction

The ability to replicate an observation by an independent study is a cornerstone of the scientific method to distinguish robust evidence from anecdote ^6^. In animal research, such replicability can be complicated by phenotypic plasticity ^3^. Whereas genotypic differences can be eliminated by selective breeding ^7–9^, the environment in which research animals are born and grow up may differ substantially between rearing facilities ^5,10,11^. As a result, genotype-by-environment interactions throughout ontogeny can lead to phenotypic differences between animals, which may hinder researchers to replicate findings, even when using genetically homogeneous (inbred) animals ^12–14^. Therefore, phenotypic plasticity may contribute to replication failure and conflicting findings in the scientific literature ^1,3,5^. However, the magnitude of this problem is as yet unknown, as existing evidence is generally based on single-laboratory studies ^15–17^ and experimentally induced environmental interventions. Here, we sought to determine the extent to which common differences in housing and husbandry conditions between rearing facilities modulate the phenotype of inbred mice, using a systematic multicenter approach.

### Study design

Pregnant C57BL/6JRj female mice from a single breeding population (Janvier Labs, Le Genest-Saint-Isle, France) were randomly allocated and transported to five independent rearing facilities (RF), where their offspring were born and reared until eight weeks of age. In order to assess the effect of the rearing environment independent of genotype and test conditions, both male and female offspring from all five RFs were then transported to a single test laboratory that was new for all mice, and after habituation period they were tested for phenotypic differences (Fig. 1a; Suppl. Fig. 1). Specifically, we examined the extent and persistence of variation in the composition of the gut microbiota associated with the different RFs and measured differences in phenotypic traits such as body weight, adrenal weight, neuroendocrine stress reactivity, and behaviour (Fig. 1b). In addition, we assessed differences in neural chromatin accessibility to explore the potential biological basis of behavioural differences (Fig. 1b). The study protocol was pre-registered (10.17590/asr.0000201) and is further detailed in the methods section.

**Figure 1.**
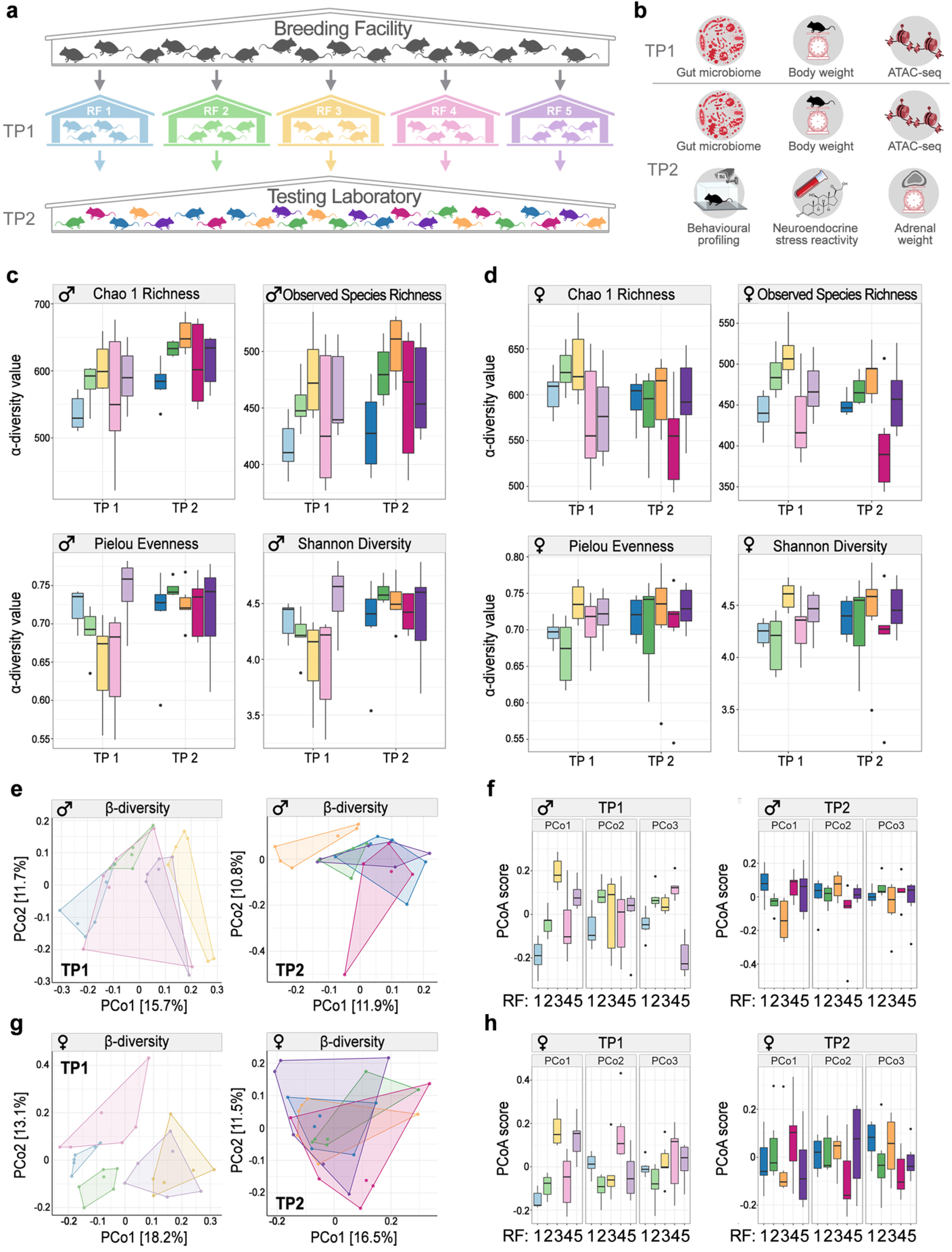
Study design and effects of rearing facility on gut microbiota diversity and composition. **a)** Schematic illustration of the multicenter study design - genetically homogeneous mice originating from a single inbred stock were reared until the age of 8 weeks in 5 different rearing facilities (RFs) before testing for phenotypic differences induced by the different rearing conditions in a single testing laboratory. **b)** Effects of the RF were evaluated at two time points (TP), first, at the end of the rearing period in each of the five RFs (TP1), and during the testing period in the testing laboratory (TP2). Outcome measures assessed at both TP1 and TP2 included gut microbiota, body weight and chromatin profiles using ATAC-seq, while behavioral tests (open field and light dark box tests) and physiological measures of stress (HPA axis reactivity test (SRT) and relative adrenal weight were limited to TP2). Values for α-diversity metrics for **c)** male mice and **d)** female mice from different rearing facilities (n = 6 mice/sex/RF/TP). **e-g)** Ordination plots visualizing Principal Coordinate analysis (PCoA) based on Bray-Curtis dissimilarity between samples of male **(e)** and female **(g)** mice from different rearing facilities, split by TPs. **f-h)** Differences between loadings of samples on the first three PCoA axes in male **(f)** and female **(h)** mice. Box plots show the first and third quartiles; horizontal line represents the median; whiskers represent the mean variability outside the upper and lower quartiles. Individual points represent outliers. TP1: 8 weeks of age PND 56; TP2: 14.5 weeks of age (PND 104). PND: postnatal day

## Results

### Rearing facility shaped the gut microbiota composition

The gut microbiome has been reported to play an important role in shaping the host phenotype ^18–22^. Therefore, we first examined the extent to which the composition of the gut microbiota varied in genetically homogeneous mice in response to the macroenvironments of the different RFs, and whether these differences persisted after the transfer to the common macroenvironment of the test laboratory.

We first analysed whether the gut microbiome of mice from different RFs differed in α-diversity measures. In males, effects of the RF were significant for both predicted and observed taxa richness, but there was little effect on overall diversity and evenness (Fig. 1c, Extended Data Table 1). At TP2, we observed an increase in all metrics of *α*-diversity, except for observed species richness (Fig. 1c; Extended Data Table 1). Similar patterns in terms of differences between RFs for taxa richness were observed in females, however, there was no change in *α*-diversity metrics across time points (Fig. 1d; Extended Data Table 1). Taken together, these results suggest that the macroenvironment of the rearing facility can lead to significant differences in the richness of the gut microbiome community.

Next, we evaluated the differences in the microbiome composition (*β*-diversity) based on the Bray-Curtis dissimilarity between samples ^23^. Overall, we found pronounced differences between mice reared in different RFs. When assessing the amount of variation in the data explained by RF at each time point, the effect was most pronounced at TP1, accounting for 28.7% of overall variation in males (Fig. 1e,f; Extended Data Table 2) and 29% in females (Fig. 1g,h; Extended Data Table 2). Importantly, the differences in microbiome composition persisted across time points, although the amount of variation explained by RF dropped to 20.4% in males and 17% in females. This implies that a large part of the initial differences in the microbiomes of mice from different RFs persisted throughout the 6 weeks in the test laboratory, although they did converge to some extent once they were housed together at the same test facility. When further investigating how overall community composition varied across RFs, we found a clear separation along principal coordinate axis 1 (PCoA1) between RFs 3 and 5 and RFs 1, 2 and 4 in both males (both time points) and females (TP1). Clustering analysis suggested that the type of mouse diet (specifically diet supplier; Suppl.Table 1a) was driving the grouping of the mice into these two populations (Suppl. Fig. 2a; Supplementary Dataset 1). Interestingly, these two populations differed in abundance of *Firmicutes* and *Bacteroidetes* (Suppl. Fig. 2b), two phyla that are associated with numerous phenotypic differences in health and disease in animal and human studies ^24–26^. Overall, these findings show that the rearing environment can lead to significant and temporally persistent compositional differences in the gut microbiome community, which in turn may drive phenotypic variation^21^.

### Rearing facility persistently affected body weight

Mice reared in different RFs differed markedly in body weight, and these differences persisted at the test laboratory throughout the experiment. RF was the only factor that had a strong and persistent effect on body weight in both males and females, while variation in litter size, litter sex ratio and group size after weaning had no significant effects (Fig. 2a; Extended Data Table 3).

**Figure 2.**
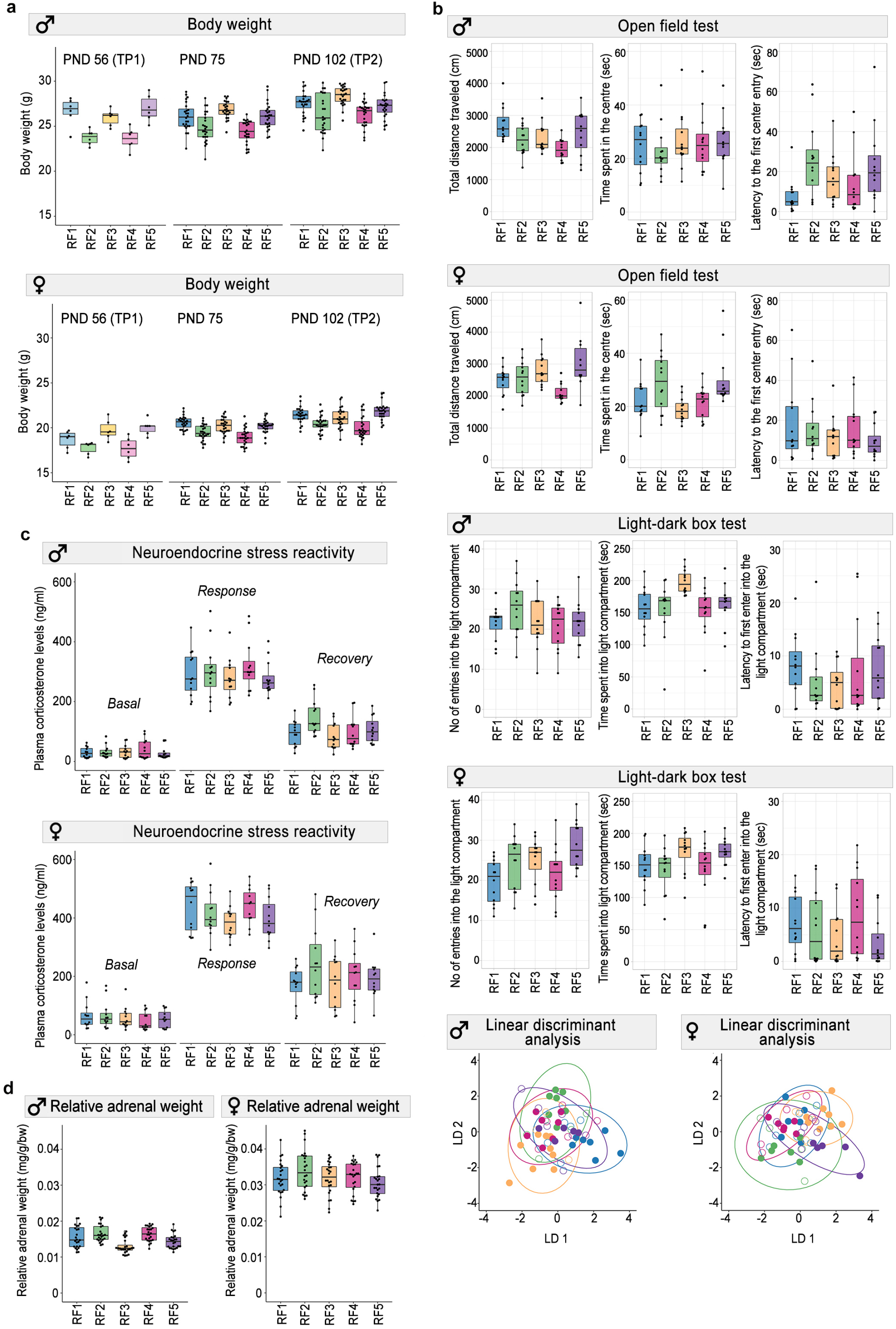
Effects of rearing facility on the behavioral and physiological profile of the mice. **a)** Body weight persistently varied by RF in both males and females (n=6 mice/sex/RF for TP1; n=24 mice/sex/RF for TP2). **b)** Behavior of the mice varied consistently by RF both in males and females. In the LDA plots, color indicates RF, and the circles represent classification based on discriminant function analysis (n=12 mice/sex/RF). **c)** RF did not affect plasma corticosterone levels in the SRT both in males and females (n=12/mice/sex/RF), while relative adrenal gland weights (n=24/mice/sex/RF) were affected only in males **(d)**. Box plots include individual data points and show the first and third quartiles; horizontal line is the median; whiskers represent the variability outside the upper and lower quartiles.

### Rearing facility modulated the behavioural phenotype

Phenotypic differences in behaviour were analysed by combining behavioral variables derived from two standard behavioural tests (open field, OF, and light-dark-box, LDB) in a multivariate analysis of variance (MANOVA). Again, we found that RF had a strong effect on the behavioural phenotype, explaining 21.6% and 18.5% of the variation in males and females, respectively. Moreover, whereas variation in litter size, litter sex ratio and group size after weaning had no effect on behaviour in both males and females, oestrous cycle stage on the test day had a strong effect in females (Extended Data Table 4; Supp. Fig. 3a-c).

Using linear discriminant function analysis (LDA) on the combined behavioural data (Extended Data Table 5), we were able to correctly classify 58% of all male mice and 53% of all female mice according to their RF, which is substantially more than the 20% predicted by chance (*χ^2^* =55.1, p=1.14×10^-13^ and *χ^2^* =41.7, p=1.08×10^-10^, respectively for males and females, Fig. 2b; Extended Data Table 6). In males, the first two discriminant functions together explained 79% of the variation between RFs, whereby the coefficients of the discriminant functions indicate that distance travelled in the OF and time in the light compartment in the LDB, two main measures of exploration and emotionality, contributed most to the first function, while time in the light compartment and number of entries into the light compartment in the LDB contributed most to the second function. In females, the first two functions together explained even 90% of the between-facility variance.

Distance travelled in the OF contributed most to the first function, while time in the centre in the OF contributed most to the second function. These findings demonstrate that common environmental differences between animal facilities can substantially alter key aspects of the behavioural phenotype of mice.

### Rearing facility did not affect neuroendocrine stress reactivity but adrenal weight

Further, we examined whether RF altered the animals’ neuroendocrine stress reactivity to a brief period of physical restraint by measuring changes in plasma corticosterone. There were no consistent differences in neuroendocrine stress reactivity between mice reared in different RFs (Fig. 1c; Extended Data Table 7). In males, there was a strong effect of litter size on basal corticosterone levels, and group size after weaning strongly affected both basal levels and acute response levels of corticosterone (Extended Data Table 6). As expected, corticosterone levels in females were almost double those in males ^27,28^ (Fig. 1g,h), and the oestrous cycle stage had a strong effect on basal corticosterone levels (Extended Data Table 7; Supp. Fig. 4). However, we found that RF had a strong effect on adrenal weight, at least in males (Fig. 1d; Extended Data Table 8). These results suggest that the chronic stress engendered by standard housing conditions and husbandry procedures induced changes in adrenal gland morphology and function, which may have buffered the neuroendocrine stress response to acute stressors ^29^.

### Rearing facility influenced chromatin organization in neuronal nuclei

We next explored whether epigenetic differences at the level of chromatin accessibility can explain some of the observed differences between mice reared in different RFs. To do so, we performed the assay for transposase-accessible chromatin using sequencing (ATAC-seq ^30^), which was applied to neuronal nuclei extracted from the ventral hippocampus, a brain area involved in modulating emotional behavior and stress responses in mice ^31,32^. This analysis was limited to males, as they showed more pronounced phenotypic variation, especially in behavioural traits.

We found that most samples clustered based on RF, suggesting pronounced differences in chromatin accessibility. This pattern was observed by looking at both the overall dissimilarity of all ATAC-seq profiles and the 10% most variable peaks (Fig. 3a). RF explained 55.33% and 36.79% of overall variation at TP1 and TP2, respectively (Fig. 3b; Extended Data Table 9; Supp. Fig. 5). Remarkably, variation explained by RF was much larger in the open chromatin sites (77.5% at TP1 and 70.9% at TP2) than in the closed sites (48.2% at TP1 and 28.4% at TP2), suggesting that these differences have functional consequences (Fig. 2b; Extended Data Table 9).

**Figure 3.**
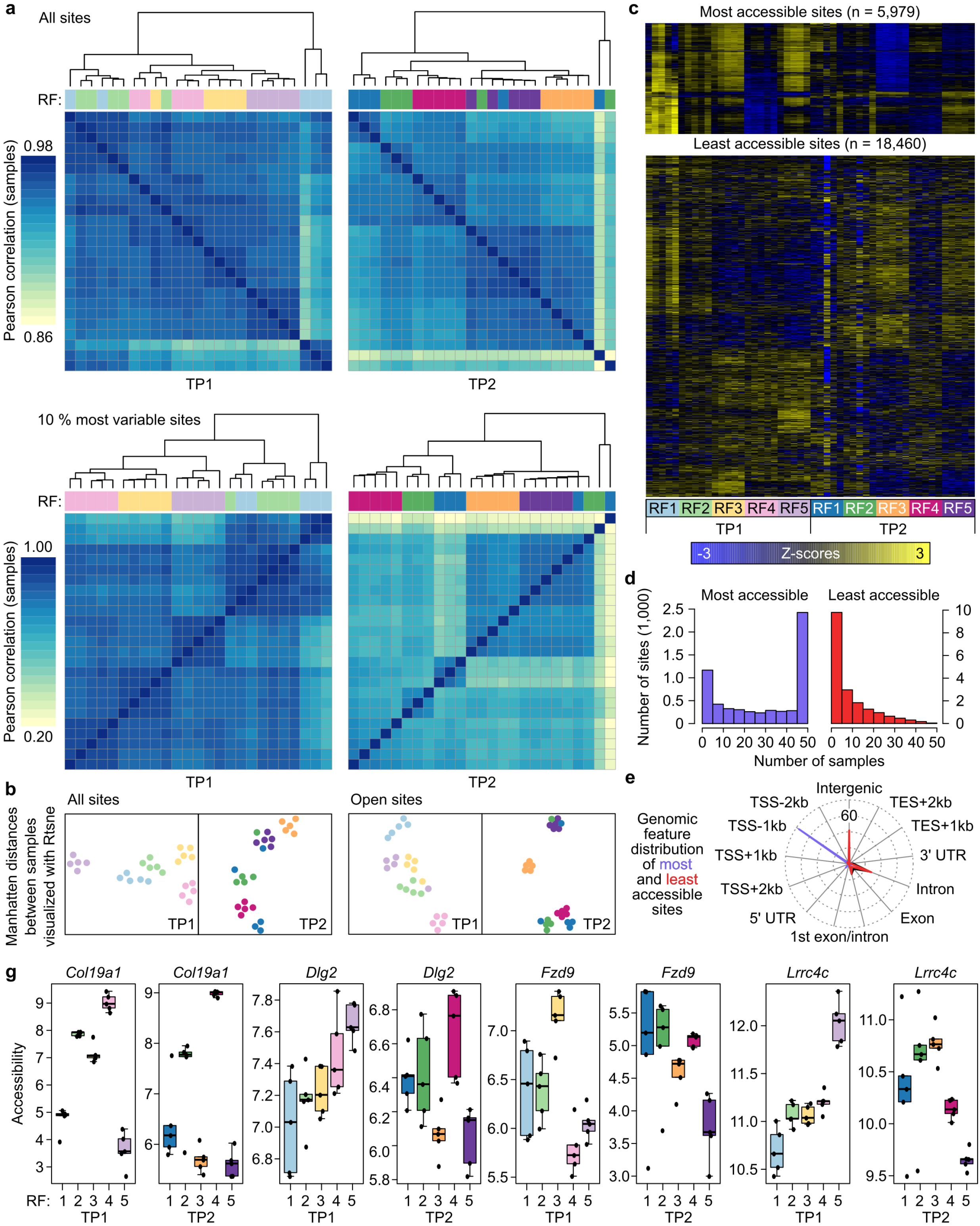
Differences in neuronal chromatin accessibility between males from different rearing facilities. **a)** Sample correlation matrices based on all sites and the 10% most variable ATAC-seq sites (n=5 mice/RF/TP). **b)** Manhatten distances between samples for all sites and open chromatin sites visualized by t-SNE. **c)** ATAC-seq accessibility signal in response to the different RFs and TPs. Heatmap representation of the most and least accessible sites. The color represents the intensity of chromatin accessibility, from gain (yellow) to loss (dark blue), calculated by using row wise Z-scores (the values are scaled by subtracting the average across samples and by dividing by the standard deviation across samples). **d)** Bar graphs representing the number of genes associated with the most and least accessible peaks. **e)** Spidergraph representing the genomic features mapped by all (black), open (blue), or closed (red) sites. **f)** Chromatin accessibility profiles of *Col19a1, Dlg 2, Fzd9* and *Lrrc4c.*

*In terms of genomic features, the* most accessible sites were preferentially located within the promoter regions, further corroborating the potential functional significance of the observed changes, while the less accessible sites were mainly located in the intergenic regions and introns (Fig. 3c, e). In addition, the number of genes associated to the most accessible peaks were common between subjects, whereas less accessible sites and associated genes behaved much more randomly and decreased with the number of selected samples (Fig. 3d).

Next, we generated lists of all differentially accessible regions (DARs) in the ventral hippocampus of mice from different RFs and mapped them to their adjacent genes for all comparisons at both TPs separately (Supplementary Data 2). Among the genes with the highest fold change, we for instance found *Col19a1* (encoding nonfibrillar collagen XIX), where chromatin accessibility differed between RFs and remained consistent across both TPs (Fig. 2g; Suppl. Fig. 5e). This indicates that persistent chromatin regulation occurred during the rearing period in response to the specific environment of the RF in genes necessary for hippocampal synapse formation ^33–35^.

We also found DARs between RFs that changed between TP1 and TP2, and mapped them to genes such as *Dlg 2* (discs large homolog 2, also known as postsynaptic density protein-93 (PSD-93), *Fzd9* (encoding Frizzled9, one of the Wnt receptors) and *Lrrc4c* (encoding Leucine Rich Repeat Containing 4C). Changes in these genes, important for postsynaptic plasticity (Fig. 2g; Suppl. Fig. 5e) ^36–39^, indicate that mice from different RFs were using different chromatin regulation strategies to adapt to the new environment of the test laboratory.

### Rearing facility induced chromatin changes relevant to neuronal function

To assess the functional significance of environmentally induced chromatin changes, we performed Gene Ontology (GO) and Kyoto Encyclopedia of Genes and Genomes (KEGG) pathway analyses, which focused on genes mapped to DARs located near transcription start site (TSS).

The GO analysis revealed that differences between RFs persistently influenced nucleosome function and regulatory processes important for hippocampal synaptic plasticity and neurogenesis, such as the response to epidermal growth factor (EGF)^40^, regulation of Notch ^41–43^, and Transforming growth factor beta (TGFβ) receptor signaling ^44,45^ (Fig. 4 a,b). There was also a clear effect of RF on genes involved in the regulation of *various behavioral processes* and presynaptic plasticity events, targeting mainly GABAergic and glutamatergic transmission (Supplementary Data 3a,c). This effect was evident only at TP1, while at TP2 enriched terms were associated with the modification of post-synaptic structure, regulation of actin cytoskeleton, dendrite development and neurotransmitter receptor complex (Supplementary Data 4b,d; Fig. 4 a,b).

**Figure 4.**
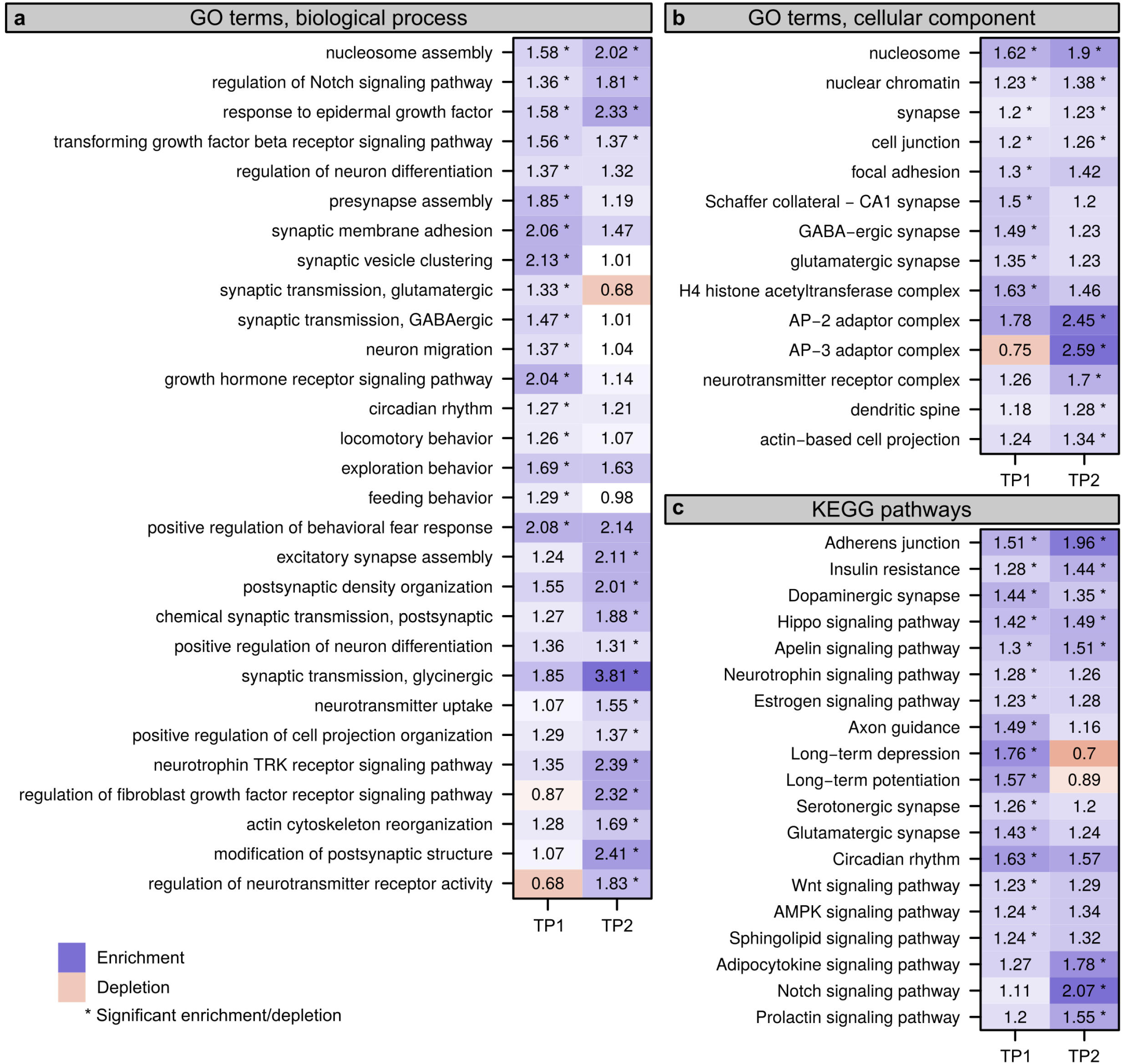
Predicted biological processes, cellular components and KEGG pathways affected by differential rearing environment. GO analysis of genes showing differential chromatin accessibility between the different RF presented for each time point (**a-b**); KEGG pathway analysis of the genes showing differential chromatin accessibility between the different rearing facilities for each TP. Fields marked with an asterisk depict comparisons that were statistically significant between TPs (two-sided Fisher’s exact test, adjusted for multiple testing, FDR < 0.05).

The KEGG analysis highlighted an overrepresentation of genes belonging to adherens junction, dopaminergic synapses, hippo, apelin and insulin resistance signaling pathway. These pathways, which have an important role in maintaining hippocampal development, morphology, and plasticity, were significantly affected by the RF at both TPs and likely have functional consequences for behavioral regulation ^46–49^. Enrichment of genes relevant to neurotrophin, long-term depression and potentiation, serotonergic and glutamatergic synapse pathways was evident only at TP1 (Fig. 4c; Supplementary Data 4a). These differences diminished after the mice had spent 6 weeks in the test laboratory. At TP2, we noticed significant differences for Notch, prolactin, relaxin, and AMPK signaling pathways, indicating their potential role in behavioral adaption to the new environment (Fig. 4c; Supplementary Data 4a).

Overall, these findings demonstrate that facility-specific macroenvironments influenced developmental programs during the late prenatal, early postnatal and adolescent period, by affecting neuronal chromatin accessibility profiles and shaping the mice’ behavioral phenotypes.

## Discussion

In this study, we found that common differences in standard housing and husbandry practices between animal facilities substantially modulated morphological, physiological and behavioural traits in mice, thereby producing facility-specific phenotypes. Our findings suggest that environmental differences between rearing facilities influence neuronal developmental patterns by modulating gene regulatory networks involved in the regulation of hippocampal synaptic plasticity and neurogenesis. Such effects on the chromatin profile of functionally relevant genes may be responsible for persistent changes in behavioral traits ^50,51^.

However, we also found that some of the pathways affected by the rearing environment maintained plasticity, possibly to facilitate adaptation to environmental change, as shown by chromatin reorganization in response to the transfer to the test laboratory at 8 weeks of age.

In conclusion, our findings could help to explain replicability issues in animal research ^52,53^. Poor replicability has mostly been attributed to publication bias, lack of statistical power, analytical flexibility and other risks of bias ^54–58^, albeit empirical evidence has remained elusive ^59^. In contrast, the large between-study heterogeneity caused by rigorous within-study standardization has long been ignored as a cause of poor replicability, despite both theoretical and empirical evidence ^1–5,60–64^. Our findings highlight an important limitation of inferences from single-laboratory studies and suggest that the (early) environmental background of animals – just like their genetic background ^65–67^ – needs to be accounted for by study design to produce robust and replicable research findings. Thus, results from standardized single-laboratory studies should be considered as preliminary evidence. Using animals from multiple breeding or rearing facilities may provide a solution to systematically heterogenize the environmental background of study populations. However, we hope that our findings will stimulate research to find other, more practicable ways to produce robust and replicable research *results.*

## Data availability

The ATAC-seq data are available from the NCBI Gene Expression Omnibus (GEO) database under accession number GSE191125. The 16S rRNA gene sequencing data is uploaded to the European Nucleotide Archive (ENA) under accession number PRJEB49361 and will be made publicly available upon acceptance of the manuscript. All other relevant data supporting the key findings of this study are available within the article and its supplementary material. A reporting summary for this article is available as a supplementary information file.

## Code availability

The code files for the ATAC-seq analysis are available at: https://github.com/MWSchmid/Jaric-et-al.-2022. The scripts used for gut microbiome, behavioral and physiological analyses are available as supplementary PDF file.

## Contributions

I.J., B.V., and H.W. designed the study; I.J. coordinated the project, performed tissue collections, testing for behavioral and physiological responses, nuclei sorting and ATAC-seq experiments; B.V performed statistical analysis of behavioral and physiological measurements with the input from I.J. and H.W. I.J., M.C. and S.S. designed the microbiota part of the study; M.C. performed DNA extraction, 16S rRNA library preparation and downstream analysis; S.S. provided the funding, contributed computational resources and supervised the microbiota part of the study. M.W.S. performed the ATAC-seq bioinformatics analysis, with the input from I.J. J.N. and M.R. assisted with behavioural experiments and stress reactivity test; R.R. helped with blood collection for the stress reactivity test; V.T.vK, M.B., I.A. and J.N. assisted with tissue collection within rearing facilities; H.R., A.B., and D.W. provided the facility resources, and laboratory space in RF 1, 2, 4 and 5. C.T. was responsible for the blood corticosterone measurements.

I.J., B.V., M.C., and H.W interpreted the data. I.J., M.C. and M.W.S. constructed the figures. H.W provided the main funding and supervised the project. I.J., B.V and H.W. wrote the manuscript with input from all authors.

## Acknowledgements

This study was supported by the Swiss National Science Foundation (grant 310030_179254) to H. W and core funding from ETH Zürich to the Laboratory of Microbiome Research headed by S.S. We would further like to acknowledge the resources of the Flow Cytometry and Cell Sorting Core Facility (FCCS CF) of the Department for BioMedical Research and the Next Generation Sequencing (NGS) Core Platform of the University of Bern. Also, we would like to thank Dr. Stefan Müller for his assistance and support with nuclei sorting as well as Dr. Pamela Nicholson for assistance with Illumina sequencing. The authors thank Nadine Sudhof for excellent technical assistance and running the plasma corticosterone assays.

## Extended Data Tables

**Extended Data Table 1:** ANOVA results for the effect of rearing facility (RF) and time point (TP) on α-diversity for males and females separately.

| Sex | Response value | Covariate | Df | SumSq | MeanSq | F | p |
| --- | --- | --- | --- | --- | --- | --- | --- |
| Males | Shannon diversity | RF | 4 | 0.6656 | 0.16639 | 1.44 | 0.2343 |
|  |  | TP | 1 | 0.6791 | 0.67908 | 5.8769 | 0.0189* |
|  |  | Residuals | 51 | 5.8931 |  |  |  |
|  | Chao1 richness | RF | 4 | 32444 | 8110.9 | 3.9072 | 0.0076* |
|  |  | TP | 1 | 25840 | 25839.7 | 12.4477 | 8.959×10<sup>-04</sup> * |
|  |  | Residuals | 51 | 105869 | 2075.9 |  |  |
|  | Observed species richness | RF | 4 | 29390 | 7347.6 | 4.6617 | 0.0028* |
|  |  | TP | 1 | 4636 | 4636 | 2.9413 | 0.0924 |
|  |  | Residuals | 51 | 80384 | 1576.2 |  |  |
|  | Pielou's evenness | RF | 4 | 0.019379 | 0.004845 | 1.8004 | 0.1431 |
|  |  | TP | 1 | 0.013766 | 0.013766 | 5.1158 | 0.0280* |
|  |  | Residuals | 51 | 0.137234 | 0.002691 |  |  |
| Females | Shannon diversity | RF | 4 | 0.7651 | 0.191279 | 1.9047 | 0.1229 |
|  |  | TP | 1 | 0.0059 | 0.005884 | 0.0586 | 0.8096 |
|  |  | Residuals | 54 | 5.4228 | 0.100423 |  |  |
|  | Chao1 richness | RF | 4 | 21502 | 5375.6 | 2.9983 | 0.0263* |
|  |  | TP | 1 | 4302 | 4301.9 | 2.3995 | 0.1272 |
|  |  | Residuals | 54 | 96815 | 1792.9 |  |  |
|  | Observed species richness | RF | 4 | 49045 | 12261.2 | 9.928 | 4.276×10<sup>-06</sup> * |
|  |  | TP | 1 | 3604 | 3603.7 | 2.918 | 0.0933 |
|  |  | Residuals | 54 | 66690 | 1235 |  |  |
|  | Pielou's evenness | RF | 4 | 0.012973 | 0.003243 | 1.4114 | 0.2426 |
|  |  | TP | 1 | 0.000714 | 0.000714 | 0.3105 | 0.5797 |
|  |  | Residuals | 54 | 0.124094 | 0.002298 |  |  |

**Extended Data Table 2:** PERMANOVA partitioning variation in microbiome microbial community composition (β-diversity) between rearing facilities (RF), for each timepoint (TP) and sex separately.

| Sex | Time point (TP) | Covariate | Df | SumSq | R2 | F | p |
| --- | --- | --- | --- | --- | --- | --- | --- |
| Males | TP1 | RF | 4 | 1.3324 | 0.2872 | 2.5179 | 9.999×10<sup>-05</sup> * |
|  |  | Residual | 25 | 3.3073 | 0.7128 |  |  |
|  |  | Total | 29 | 4.6397 | 1 |  |  |
|  | TP2 | RF | 4 | 0.7666 | 0.2038 | 1.4079 | 0.0004 * |
|  |  | Residual | 22 | 2.9947 | 0.7962 |  |  |
|  |  | Total | 26 | 3.7613 | 1 |  |  |
| Females | TP1 | RF | 4 | 1.107 | 0.2908 | 2.5628 | 9.999×10<sup>-05</sup> * |
|  |  | Residual | 25 | 2.6996 | 0.7092 |  |  |
|  |  | Total | 29 | 3.8066 | 1 |  |  |
|  | TP2 | RF | 4 | 0.6665 | 0.1699 | 1.2791 | 0.0444 * |
|  |  | Residual | 25 | 3.2567 | 0.8301 |  |  |
|  |  | Total | 29 | 3.9231 | 1 |  |  |

**Extended Data Table 3:** Phenotypic variation in body weight of mice is induced by common differences between the rearing conditions in different facilities. The effect of rearing environment on body weight was evaluated at three time points throughout the study: right before euthanasia within each rearing facility (PND 56; TP1), after the acclimatization period in the testing facility (PND 75) and at the end of the experiment (PND 102; TP2). Linear models were used to analyze data collected on mouse body weight at PND 56 (TP1) within each rearing facility for males and females (a). Rearing facility, litter size and sex ratio at weaning were used as predictor variables. Linear mixed effect models with the same list of predictor variables as fixed effects were used for the body weight data collected in the testing facility for males and females (b). Cage identification number (cage ID) in the testing facility was used as a random factor.

| Sex | Variables | Df | Sum Sq | Mean Sq | F value | p |
| --- | --- | --- | --- | --- | --- | --- |
| Males | Rearing Facility | 4 | 62.619 | 15.6525 | 9.7496 | $9.185 \times 10^{-05} *$ |
|  | Litter size at weaning | 1 | 0.577 | 0.5769 | 0.3593 | 0.5547<sup>ns</sup> |
|  | Sex ratio at weaning | 1 | 0.014 | 0.0141 | 0.0088 | 0.9263<sup>ns</sup> |
|  | Residuals | 23 | 36.925 | 1.6055 |  |  |
| Females | Rearing Facility | 4 | 28.6847 | 7.1712 | 8.4225 | $2.447 \times 10^{-04} *$ |
|  | Litter size at weaning | 1 | 4.1711 | 4.1411 | 4.8989 | 0.0371<sup>ns</sup> |
|  | Sex ratio at weaning | 1 | 0.371 | 0.371 | 0.4357 | 0.5158<sup>ns</sup> |
|  | Residuals | 23 | 19.583 | 0.8514 |  |  |

| Sex | PND | Fixed factors | Sum Sq | Mean Sq | num Df | den Df | F value | p |
| --- | --- | --- | --- | --- | --- | --- | --- | --- |
| Males | BW after acclimatization period (PND 75) | Rearing Facility | 20.5856 | 5.1464 | 4 | 52 | 5.1178 | 0.0015* |
|  |  | Litter size at weaning | 1.0737 | 1.0737 | 1 | 52 | 1.0677 | 0.3062<sup>ns</sup> |
|  |  | Sex ratio at weaning | 0.5034 | 0.5034 | 1 | 52 | 0.5006 | 0.4824<sup>ns</sup> |
|  |  | Number of cage mates after weaning | 0.1953 | 0.1953 | 1 | 52 | 0.1943 | 0.6612<sup>ns</sup> |
|  |  | Random factor: Cage ID in the testing lab |  |  |  |  |  |  |
|  |  | REML criterion at convergence: 399.7 |  |  |  |  |  |  |
|  |  | marginal R2 0.3071549; conditional R2 0.6517345 |  |  |  |  |  |  |
|  | BW at the end of the experiment (PND 102; TP2) | **Fixed factors** | **Sum Sq** | **Mean Sq** | **num Df** | **den Df** | **F value** | **p** |
|  |  | Rearing Facility | 19.1941 | 4.7985 | 4 | 52 | 3.5935 | 0.0116* |
|  |  | Litter size at weaning | 0.2587 | 0.2587 | 1 | 52 | 0.1938 | 0.6616<sup>ns</sup> |
|  |  | Sex ratio at weaning | 0.8734 | 0.8734 | 1 | 52 | 0.6541 | 0.4223<sup>ns</sup> |
|  |  | Number of cage mates after weaning | 1.0087 | 1.0087 | 1 | 52 | 0.7554 | 0.3888<sup>ns</sup> |
|  |  | Random factor: Cage ID in the testing lab |  |  |  |  |  |  |
|  |  | REML criterion at convergence: 429.4 |  |  |  |  |  |  |
|  |  | marginal R2 0.2106322; conditional R2 0.5912489 |  |  |  |  |  |  |
| Females | BW after acclimatization period (PND 75) | **Fixed factors** | **Sum Sq** | **Mean Sq** | **num Df** | **den Df** | **F value** | **p** |
|  |  | Rearing Facility | 18.1681 | 4.542 | 4 | 52 | 9.0795 | 1.221×10<sup>-05</sup>* |
|  |  | Litter size at weaning | 0.1548 | 0.1548 | 1 | 52 | 0.3095 | 0.5804<sup>ns</sup> |
|  |  | Sex ratio at weaning | 0.074 | 0.074 | 1 | 52 | 0.1479 | 0.7022<sup>ns</sup> |
|  |  | Number of cage mates after weaning | 0.0361 | 0.0361 | 1 | 52 | 0.0722 | 0.7892<sup>ns</sup> |
|  |  | Random factor: Cage ID in the testing lab |  |  |  |  |  |  |
|  |  | REML criterion at convergence: 399.7 |  |  |  |  |  |  |
|  |  | marginal R2 0.3071549; conditional R2 0.6517345 |  |  |  |  |  |  |
|  | BW at the end of the experiment (PND 102; TP2) | **Fixed factors** | **Sum Sq** | **Mean Sq** | **num Df** | **den Df** | **F value** | **p** |
|  |  | Rearing Facility | 17.5852 | 4.3963 | 4 | 52 | 6.9142 | 1.533×10<sup>-04</sup>* |
|  |  | Litter size at weaning | 0.0314 | 0.0314 | 1 | 52 | 0.0494 | 0.8250<sup>ns</sup> |
|  |  | Sex ratio at weaning | 0.0118 | 0.0118 | 1 | 52 | 0.0186 | 0.8921<sup>ns</sup> |
|  |  | Number of cage mates after weaning | 0.2106 | 0.2106 | 1 | 52 | 0.3312 | 0.5674<sup>ns</sup> |
|  |  | Random factor: Cage ID in the testing lab |  |  |  |  |  |  |
|  |  | REML criterion at convergence: 429.4 |  |  |  |  |  |  |
|  |  | marginal R2 0.2106322; conditional R2 0.5912489 |  |  |  |  |  |  |

**Extended Data Table 4:**
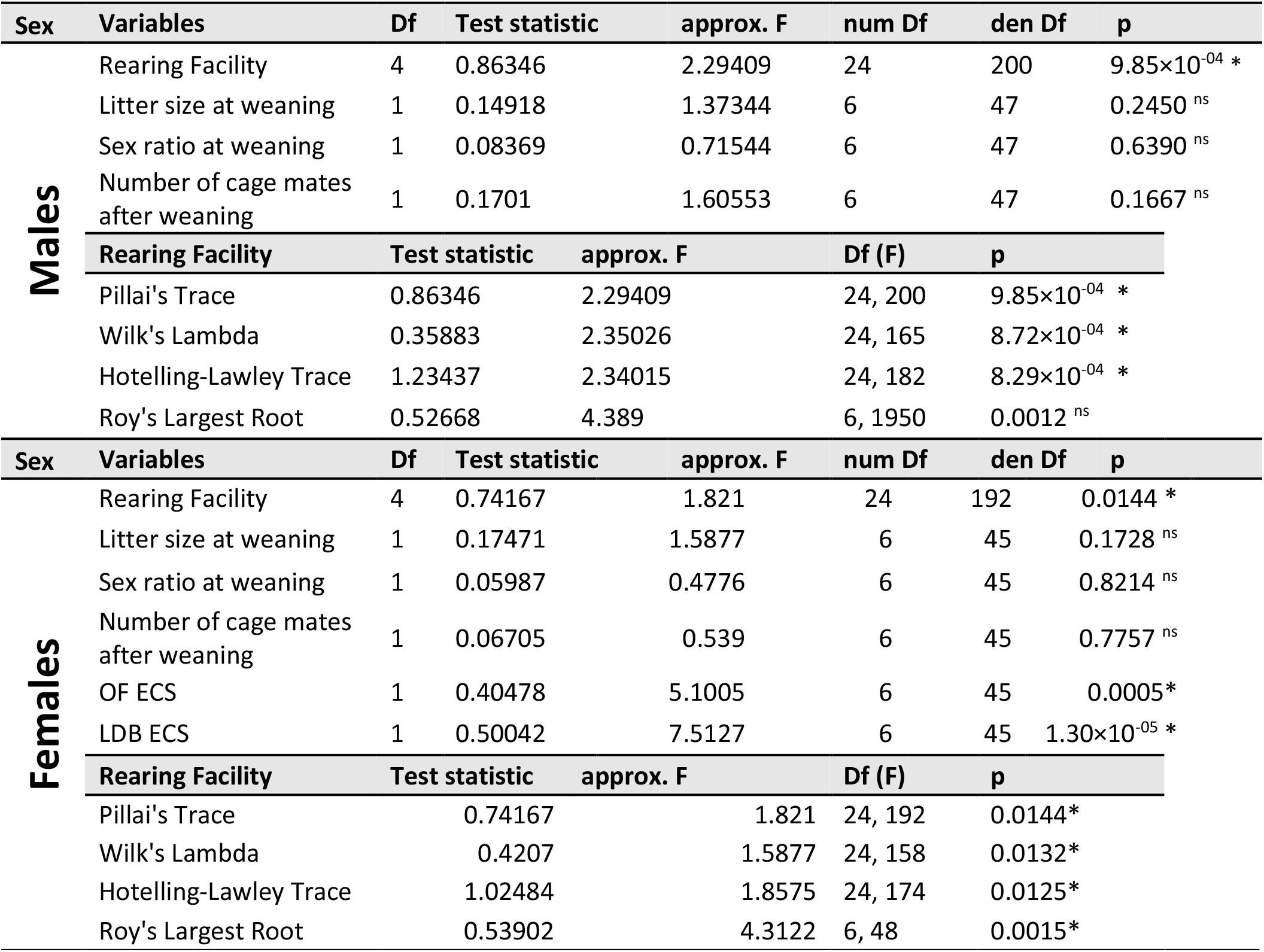
Phenotypic variation in behavior of mice is modified by common differences between the rearing conditions in different facilities. MANOVA outcomes and statistics for males and females. The rearing facility was defined as a main independent (predictor) variable. Additional covariates were included in the model based on the published evidence, which suggests that they might affect behavioral phenotype. List of the nuisance variables includes: litter size at weaning ^93,94^, sex ratio at weaning ^95,96^ and number of cage mates after weaning ^97^. In addition, we included the stage of oestrous cycle at the time of testing (OF ECS and LDB ECS) in females, because of its effects on behaviour ^78,80,98^. The results were confirmed by comparing the Pillai’s Trace outcome with outcomes of three different test statistics. The proportion of variation in behaviour of mice, which is solely attributed to differences in rearing environments, was calculated by dividing the Pillai’s trace by the degrees of freedom. MANOVA outcomes and statistics for behavioural parameters.

**Extended Data Table 5:**
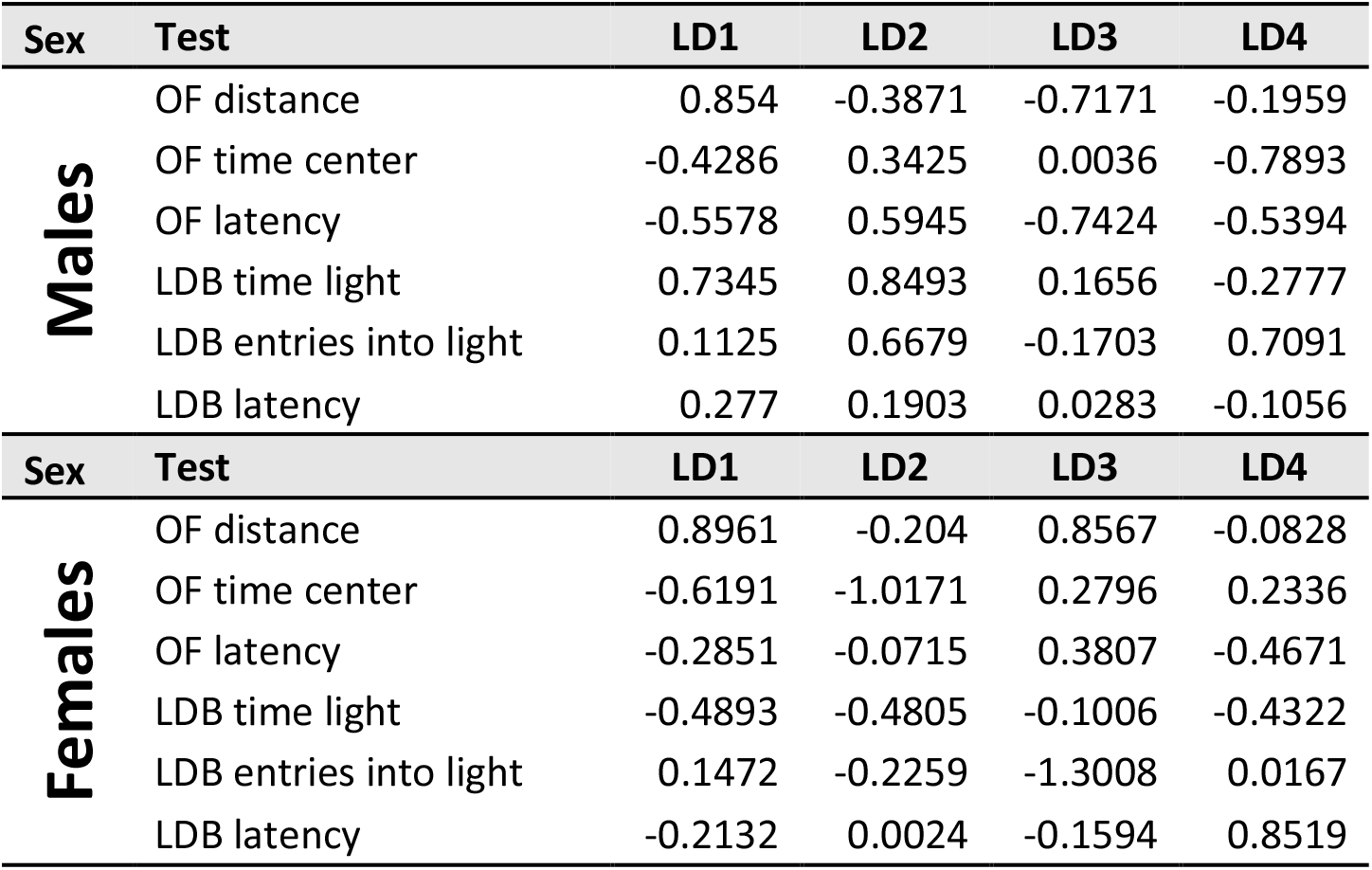
Loadings of a LDA for male and female behaviour. The rearing facility was defined as a grouping variable, while the six behavioral measures served as predictor variables. (OF=open field test, LDB=light-dark box test).

| Sex | Test | LD1 | LD2 | LD3 | LD4 |
| --- | --- | --- | --- | --- | --- |
| **Males** | OF distance | 0.854 | -0.3871 | -0.7171 | -0.1959 |
|  | OF time center | -0.4286 | 0.3425 | 0.0036 | -0.7893 |
|  | OF latency | -0.5578 | 0.5945 | -0.7424 | -0.5394 |
|  | LDB time light | 0.7345 | 0.8493 | 0.1656 | -0.2777 |
|  | LDB entries into light | 0.1125 | 0.6679 | -0.1703 | 0.7091 |
|  | LDB latency | 0.277 | 0.1903 | 0.0283 | -0.1056 |
| Sex | Test | LD1 | LD2 | LD3 | LD4 |
| **Females** | OF distance | 0.8961 | -0.204 | 0.8567 | -0.0828 |
|  | OF time center | -0.6191 | -1.0171 | 0.2796 | 0.2336 |
|  | OF latency | -0.2851 | -0.0715 | 0.3807 | -0.4671 |
|  | LDB time light | -0.4893 | -0.4805 | -0.1006 | -0.4322 |
|  | LDB entries into light | 0.1472 | -0.2259 | -1.3008 | 0.0167 |
|  | LDB latency | -0.2132 | 0.0024 | -0.1594 | 0.8519 |

**Extended Data Table 6:**
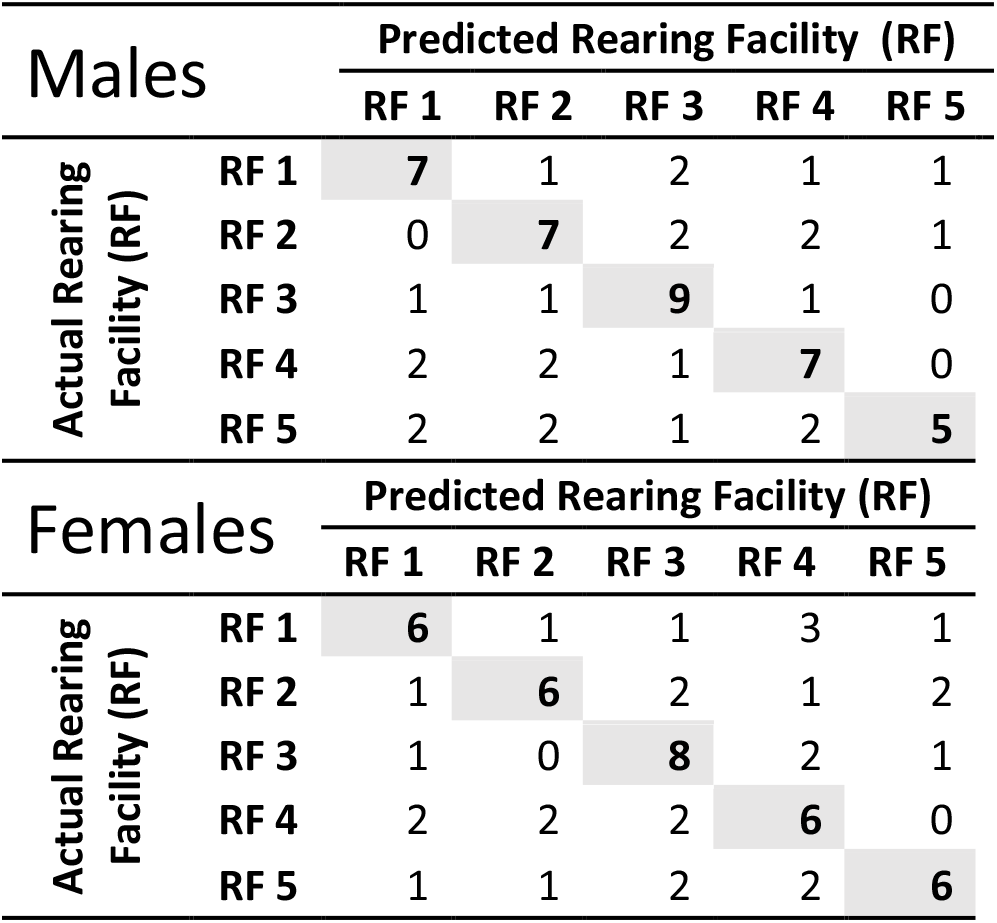
Classification based on LDA of behaviour of males and females.

**Extended Data Table 7:** Phenotypic differences of the HPA stress profile in mice cannot be explained by common differences between the rearing conditions in different facilities. Statistical outcomes of plasma corticosterone measures for males and females. We applied linear models, with the covariates rearing facility, litter size at weaning, sex ratio at weaning and number of cage mates after weaning. In females, the stage of oestrous cycle on the testing day was also included, and data were grouped into high- and low-oestrogen state.

| Sex | Value | Variables | Df | Sum Sq | Mean Sq | F value | p |
| --- | --- | --- | --- | --- | --- | --- | --- |
| Males | Basal | Rearing Facility | 4 | 1089.3 | 272.3 | 0.7134 | 0.5865<sup>ns</sup> |
|  |  | Litter size at weaning | 1 | 2820.5 | 2820.5 | 7.3891 | 0.0089* |
|  |  | Sex ratio at weaning | 1 | 83.2 | 83.2 | 0.2179 | 0.6426<sup>ns</sup> |
|  |  | Number of cage mates after weaning | 1 | 8152.7 | 8152.7 | 21.3581 | 2.549×10<sup>-05</sup>* |
|  |  | Residuals | 52 | 19849.2 | 381.7 |  |  |
|  | Response | Rearing Facility | 4 | 15273 | 3818 | 0.6879 | 0.6036<sup>ns</sup> |
|  |  | Litter size at weaning | 1 | 475 | 475 | 0.0856 | 0.7710<sup>ns</sup> |
|  |  | Sex ratio at weaning | 1 | 598 | 598 | 0.1078 | 0.7440<sup>ns</sup> |
|  |  | Number of cage mates after weaning | 1 | 39812 | 39812 | 7.1723 | 0.0099* |
|  |  | Residuals | 52 | 288646 | 5551 |  |  |
|  | Recovery | Rearing Facility | 4 | 29750 | 7437.6 | 2.9757 | 0.0275<sup>ns</sup> |
|  |  | Litter size at weaning | 1 | 27 | 27.4 | 0.011 | 0.9170<sup>ns</sup> |
|  |  | Sex ratio at weaning | 1 | 1137 | 1136.8 | 0.4548 | 0.5030<sup>ns</sup> |
|  |  | Number of cage mates after weaning | 1 | 1277 | 1276.8 | 0.5109 | 0.4780<sup>ns</sup> |
|  |  | Residuals | 52 | 129971 | 2499.4 |  |  |
| Sex | Value | Variables | Df | Sum Sq | Mean Sq | F value | p |
| Females | Basal | Rearing Facility | 4 | 3128 | 782 | 0.6902 | 0.6021<sup>ns</sup> |
|  |  | Litter size at weaning | 1 | 1296 | 1295.5 | 1.1433 | 0.2900<sup>ns</sup> |
|  |  | Sex ratio at weaning | 1 | 46 | 46.1 | 0.0407 | 0.8409<sup>ns</sup> |
|  |  | Number of cage mates after weaning | 1 | 73 | 73 | 0.0644 | 0.8006 |
|  |  | CORT ESC | 1 | 26958 | 26957.9 | 23.7911 | 1.093×10<sup>-05</sup>* |
|  |  | Residuals | 51 | 57788 | 1131.1 |  |  |
|  | Response | Rearing Facility | 4 | 39010 | 9752.5 | 1.7054 | 0.1632<sup>ns</sup> |
|  |  | Litter size at weaning | 1 | 17 | 17.5 | 0.0031 | 0.9561<sup>ns</sup> |
|  |  | Sex ratio at weaning | 1 | 471 | 471.2 | 0.0824 | 0.7752<sup>ns</sup> |
|  |  | Number of cage mates after weaning | 1 | 4331 | 4330.6 | 0.7573 | 0.3883<sup>ns</sup> |
|  |  | CORT ESC | 1 | 1542 | 1541.9 | 0.2696 | 0.6058<sup>ns</sup> |
|  |  | Residuals | 51 | 288646 | 5551 |  |  |
|  | Recovery | Rearing Facility | 4 | 41128 | 10281.9 | 1.2577 | 0.2987<sup>ns</sup> |
|  |  | Litter size at weaning | 1 | 18892 | 18892 | 2.3109 | 0.1346<sup>ns</sup> |
|  |  | Sex ratio at weaning | 1 | 1943 | 1942.9 | 0.2377 | 0.6280<sup>ns</sup> |
|  |  | Number of cage mates after weaning | 1 | 2 | 1.6 | 0.0002 | 0.9891<sup>ns</sup> |
|  |  | CORT ESC | 1 | 7510 | 7510.1 | 0.9186 | 0.3424<sup>ns</sup> |
|  |  | Residuals | 51 | 416938 | 8175.3 |  |  |

**Extended Data Table 8:** Phenotypic variation in relative adrenal weight of mice is induced by common differences between the rearing conditions in different facilities. Statistical outcomes for relative adrenal gland weight data collected at the testing laboratory (TP1; PND 56) for males and females. Linear mixed effect models were used with rearing facility, litter size at weaning, sex ratio at weaning and number of cage mates after weaning as fixed effect, while the cage ID in the testing facility was defined as a random factor. Linear mixed effect model with type III ANOVA with Satterthwaite’s approximation for relative adrenal gland weight for males.

| Sex | Fixed factors | Sum Sq | Mean Sq | num Df | den Df | F value | p |
| --- | --- | --- | --- | --- | --- | --- | --- |
| **Males** | Rearing Facility | 6.06e<sup>-05</sup> | 1.5138×10<sup>-05</sup> | 4 | 52 | 5.4086 | 1.02×10<sup>-03</sup> * |
|  | Litter size at weaning | 2.48e<sup>-06</sup> | 2.4788×10<sup>-06</sup> | 1 | 52 | 0.8857 | 0.3510 <sup>ns</sup> |
|  | Sex ratio at weaning | 1.26e<sup>-06</sup> | 1.2639×10<sup>-06</sup> | 1 | 52 | 0.4516 | 0.5045 <sup>ns</sup> |
|  | Number of cage mates after weaning | 9.61e<sup>-07</sup> | 9.6140×10<sup>-07</sup> | 1 | 52 | 0.3435 | 0.5603 <sup>ns</sup> |
|  | Random factor: Cage ID in the testing lab |  |  |  |  |  |  |
|  | REML criterion at convergence: -1032.6<br>marginal R2 0.2589189; conditional R2 0.3630702 |  |  |  |  |  |  |
| **Females** | Rearing Facility | 5.56e<sup>-05</sup> | 1.39e<sup>-05</sup> | 4 | 52 | 0.9054 | 0.4677* |
|  | Litter size at weaning | 6.99e<sup>-07</sup> | 6.99e<sup>-06</sup> | 1 | 52 | 0.0456 | 0.8318 <sup>ns</sup> |
|  | Sex ratio at weaning | 3.50e<sup>-05</sup> | 3.50e<sup>-06</sup> | 1 | 52 | 2.2827 | 0.1369 <sup>ns</sup> |
|  | Number of cage mates after weaning | 1.02e<sup>-05</sup> | 1.02e<sup>-05</sup> | 1 | 52 | 0.6636 | 0.4190 <sup>ns</sup> |
|  | Random factor: Cage ID in the testing lab |  |  |  |  |  |  |
|  | REML criterion at convergence: -1032.6<br>marginal R2 0.2589189; conditional R2 0.3630702 |  |  |  |  |  |  |

**Extended Data Table 9:** PERMANOVA partitioning variation in chromatin accessibility profile between rearing facilities and processing batches. Manhattan distance between all samples was used as input.

| Chromatin accessibility profile | Rearing Laboratory |  | Processing Batch |  |
| --- | --- | --- | --- | --- |
|  | Sum Sq | p | Sum Sq | p |
| all sites TP1 | 55.33 | 1.00e<sup>-04</sup>* | 11.13 | 0.1431 |
| all sites TP2 | 36.79 | 1.00e<sup>-04</sup>* | 12.93 | 0.3862 |
| open chromatin sites TP1 | 77.53 | 1.00e<sup>-04</sup>* | 10.01 | 0.0063 |
| open chromatin sites TP2 | 70.91 | 1.00e<sup>-04</sup>* | 6.5 | 0.3343 |
| closed chromatin sites TP1 | 48.24 | 1.00e<sup>-04</sup>* | 11.83 | 0.1998 |
| closed chromatin sites TP2 | 28.44 | 1.00e<sup>-04</sup>* | 14.37 | 0.4184 |

## Methods

### Study preregistration

Before data acquisition started, the study protocol was preregistered under the DOI number: 10.17590/asr.0000201 at the Animal Study Registry, operated by the German Centre for the Protection of Laboratory Animals (Bf3R) at the German Federal Institute for Risk Assessment (BfR).

All animal experiments were conducted in full compliance with the Swiss Animal Welfare Ordinance (TSchV 455.1) and were approved by the Cantonal Veterinary Office in Bern, Switzerland (permit number: BE12/19). Rearing facilities (RFs) located in Hannover and Münster did not require separate approval by governmental authorities for an experimental animal study, because animals were only housed in those laboratories, while all experimental procedures were carried out in Bern under the above-mentioned license.

### Animal subjects and study design

Nine-week-old, time-mated, primiparous pregnant C57BL/6JRj females in the last third of pregnancy (gestational day 14 or 15), all derived from the same breeding stock of a commercial breeder (Janvier Labs, Le Genest-Saint-Isle, France), were randomly allocated to 5 different rearing animal facilities (n = 18 per facility). The RFs were located at the following institutions: **i)** Institute of Laboratory Animal Science, Hannover Medical School, Germany (RF 1 and RF2); **ii)** Division of Animal Welfare, Vetsuisse Faculty, University of Bern, Switzerland (RF 3); **iii)** Department of Behavioural Biology, University of Münster, Germany (RF 4) and **iv)** Institute of Anatomy, University of Zürich, Switzerland (RF 5).

Pregnant dams were singly housed for approximately 5 days, from arrival at the RF until parturition. Dams were monitored daily for parturition and day of birth was defined as postnatal day 0 (PND 0). Litters were not culled during the lactation period and all healthy pups were weaned at PND 22. At weaning, in each RF 12 litters with at least 3 pups of each sex were selected randomly from all litters. If necessary, to achieve n=12, these were complemented by litters with at least 2 pups of each sex. From each litter, 3 (or 2) pups per sex were selected randomly and reared together until the age of 8 weeks (PND 56) according to the specific protocols of housing and husbandry of each of the 5 RFs (e.g. type of cages, handling method, bedding, nesting material, diet, light regime). Detailed housing and husbandry conditions are reported in Suppl. Table 1a. The first 8 weeks of postnatal life were selected because they cover two sensitive developmental periods (*i.e.* early life and adolescence) in mice ^1^. These periods represent a critical stage of brain ^2–6^, HPA axis ^7,8^ and gut microbiome development ^9,10^ when environmental inputs may shape the later-life phenotype of mice at different levels of organization.

The effect of rearing environment was evaluated at two time points (Fig. 2b; Suppl. Fig 1). At TP1, one mouse per sex and cage of all cages with 3 mice was weighed and sacrificed within each of the five RFs and brain and cecal samples were harvested for epigenomic and microbiome analyses. To ensure that observed differences are attributed only to the differential rearing environments, we standardized the tissue collection procedure. All mice were sacrificed at the beginning of the light cycle (within the first 3 hours after the lights were turned on) by the same person who was not in contact with those animals before and who was using the same equipment in each of the 5 RFs. Blinding with regard to RF was not possible for weighing and organ collection since the experimenter needed to travel to each RF.

At PND 58 the remaining pairs of male and female offspring (n = 240; 24 mice per sex per rearing) were transferred from the 5 RFs to the testing laboratory, located at the Vetsuisse Faculty of the University of Bern (testing laboratory). The testing lab in Bern was separate from the RF in Bern. To ensure that all animals experienced approximately the same treatment during transport to the testing facility, the driver of the car transporting the animals from the RF in Bern to the adjacent test laboratory in Bern was asked to take a 2-hours detour. Special efforts were made to reduce any possible transport-induced stress. The same-sex cage-mates were placed into 4 compartment transport boxes (Type 500005L; Bio Services BV, Uden, The Netherlands), with 2 mice per compartment and shipped by a professional company using an environmentally controlled vehicle. Each compartment contained 1 cm of bedding, nesting material from the home cage, food pellets and hydrogel.

Upon arrival, animals were checked for health and pair housed in freshly bedded Type 3 cages (floor area 820 cm^2^) and habituated to the new animal facility for 2.5 weeks. Each Type 3 cage contained 3 cm of bedding (OSafe® Premium Bedding, SAFE FS 14, Safe-Lab, Rosenberg, Germany), a red mouse house (Tecniplast, Indulab, Gams, Switzerland), a medium-size cardboard tunnel (Play tunnel, #CPTUN00016P, Plexx B.V. Netherlands), and 10 g of nesting material (Sizzle Nest #SIZNEST00016P, Plexx B.V. Netherlands). Standard rodent chow (Kliba Nafag #3430, Switzerland) and tap water were available ad libitum. Females and males were housed in separate rooms and all animals were kept on a 12:12 light/dark cycle, with lights on at 12:00h. Detailed housing and husbandry conditions in the testing facility are reported in Suppl. Table 1b.

The day after arrival in the testing facility, animals were individually marked by ear tattoo, after which cages were assigned new identification numbers, and positions of cages within and between cage racks were randomly re-shuffled as part of the blinding procedure. Further information on blinding is available in the reporting summary.

Behavioural and physiological phenotyping commenced after an acclimatization period of 2.5 weeks. We focused on phenotypic traits of exploration, emotionality and stress reactivity that are known to be sensitive to environmental variation during early ontogeny ^3,7,11,12^. Two common tests for exploration and emotionality, the open-field test (OF) and the light-dark box test (LDB), were conducted in that order, with a break of 7 days in between, followed by a stress reactivity test (SRT) after another break of 7 days. For these tests, one mouse per cage, sex, and RF (n=120) was used. After one additional week (around 14.5 weeks of age; PND 102; TP2), all mice (n=240) were euthanized for post-mortem analyses. Body weights were also recorded for each mouse during cage changes throughout the acclimatization period and prior to euthanasia. To avoid possible influences of the circadian rhythm on behaviour, corticosterone secretion, and molecular readouts, all procedures were performed during the light phase (from 12:00h to 16:30h).

### Tissue sampling procedure

All animals were euthanized by cervical dislocation. Whole brains were immediately removed, quickly frozen in a hexane bath on dry ice before being stored at −80°C. The brain region of interest, *i.e.* ventral hippocampus, was dissected later for subsequent molecular analyses. Adrenal glands were removed, dissected from fat, and weighed using a precision scale (Mettler AE160, Mettler-Toledo, Switzerland). The mouse cecum together with its content was isolated, snap frozen in liquid nitrogen and kept at −80°C for microbial DNA extraction and sequencing.

### Gut Microbiota Composition Analysis

For gut microbiota analysis, samples of mouse caeca were taken at both time points, at each RF and at the end of the experiment at the testing laboratory. Six cages per sex and RF were randomly selected from all cages contained 3 littermates after weaning (n=180). One mouse from each selected cage was sacrificed in the RF at 8 weeks of age (TP1; PND 56; n=60), while the other two cage mates (n=120; one underwent behavioral testing and one remained naïve) were sacrificed in the testing laboratory at TP2 (14.5 weeks of age; PND 102).

Microbial DNA was extracted from cecal content using the Allprep DNA/RNA mini kit (Qiagen, Hilden, Germany) according to the manufacturer’s instruction. DNA was eluted and concentrations were assessed using the QuantiFluor® dsDNA system (Promega).

The 16S rRNA V4 region was amplified using the following primers 515f-Y: GTGYCAGCMGCCGCGGTA and 806r-N: GGACTACNVGGGTWTCTAAT and The Q5® high-fidelity DNA polymerase kit (New England BioLabs, UK). 2μl of extracted DNA was added to a PCR reaction mix prepared by mixing a final concentration of 1X Q5 reaction buffer, 200 μM of dNTPs, 0.5 μM of each primer, 0.02 U/μL of Q5 5 High-Fidelity DNA Polymerase, and 1X of Q5 High GC enhancer, in a total volume of 25 μL. The first PCR reaction was carried out using the following conditions: i) first denaturation: 95°C for 30 sec; ii) denaturation in each PCR cycle: 98°C for 10 sec; iii) annealing: 56°C for 30 sec; iv) extension: 72°C for 30 sec; v) final extension at the end of the reaction: 72°C for 2 min, followed by a hold step at 4°C. The cycles 2-4 were repeated 8 times. The PCR products were purified using CleanNA CleanNGS purification beads (CNGS0050; LabGene Scientific SA), resuspended in 15 μl of EB buffer and served as templates in the second PCR reaction. In the second PCR step, unique dual index barcodes of length 2×8nt where added to each sample, which allowed equimolar pooling of samples after quantification of the target product using the Agilent fragment analyzer (Agilent). In total, the final library pool contained 176 samples, 3 bacterial mock communities and 20 DNA extraction blanks. The finished library pool was sequenced using the NovaSeq 6000 platform (Illumina, USA) in a single lane of SP flow cell at the Functional Genomics Center Zürich.

The raw sequencing data were analyzed using the DADA2 pipeline (version 1.14 16) and individual reads were grouped into ASVs (Amplicon sequence variants). The final table contained 1560 ASVs. After removal of the blank and mock samples from the data, the individual library sizes ranged from 47910 to 1616753, with a median of 843361 (Suppl. Fig. 6a). To mitigate the effect of variation in library size across samples, we performed random down-sampling of reads within each sample to an even library size across samples. Given the minimum read count in the data, counts were rarefied to a depth of 47,000 reads per sample (Suppl. Fig. 6b). We calculated both α diversity (within sample diversity) and β diversity (between-sample diversity) in order to assess the effect of RF on microbial community composition within and between mice.

For α diversity, the following metrics were calculated: i) Observed species richness, which represents the total number of species counted within a sample; ii) Chao1 richness, for estimation of the “true” species diversity, which is calculated using the following formula 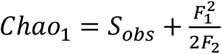, where *S_obs_* stands for the observed number of species, and *F*_1_ and *F*_2_ stand for the number of species with one or two observed reads, respectively; iii) Shannon diversity index, which illustrates the diversity within a sample, taking both richness and evenness into account. It is calculated using the following formula 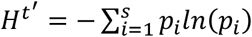, where *S_j_* represents the total number of species and *p* represents the proportion (n/N) of individuals of one particular species found (n), divided by the total number of individuals found (N), and iv) Pielou evenness for estimation of how similar in numbers each species in a sample is. It is calculated using the formula 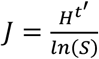, where *H^t′^* is the Shannon index and *S* is the total number of species in a sample. ANOVA was used to test the effect of RF and time point on α-diversity measures.

To analyze changes in microbiota composition between mice reared at different time points or RFs, the Bray-Curtis dissimilarity between samples was calculated using the formula 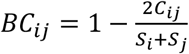, where *i* and *j* are the two samples, 5, is the total number of species counted in sample *i, S_j_* is the total number of species counted in sample site *j,* and *C_ij_* is the sum of only the lesser counts for each species found in both samples. Principal coordinate analysis (PCoA) was used to visualize the similarity in microbiome composition between samples and to retrieve sample loadings onto the first three PCoA axes. A permutational analysis of variance (PERMANOVA) was used to assess the proportion of variation in microbiome composition explained by RF and age (i.e. time point).

Based on our study design, there were three groups of cecal samples: i) samples collected at TP1 (within each of RFs); ii) samples collected at TP2 from mice that underwent behavioral testing and iii) samples collected at TP2 from mice that did not undergo behavioral testing and that were used for chromatin profiling. To balance the study design (there were double number of samples collected at TP2 comparing to the number of samples collected at TP1), we assessed whether the two sets of samples (from behaviorally tested-BT mice and mice used for molecular analysis, i.e chromatin profiling-MA mice) collected at TP2 differ in terms of species richness, diversity, and overall community composition. Our analysis showed that there were no statistically significant differences in either α diversity (assessed using Wilcoxon signed-rank test) or β diversity (assessed using PERMANOVA) in any of the tested parameters between BT and MA mice (Suppl. Fig. 7). Therefore, only MA-mice were considered for the final analyses. Differential abundance analysis was performed by using an adjusted p-value threshold of 0.05 and a log2-foldchange threshold of 1.

All analyses related to the gut microbiome were done in R (version 4.1.0) using the libraries vegan (2.5-7) for down sampling, PERMANOVA and calculation of observed/chao1 species richness, microbiome (1.14.0) for calculation of Shannon diversity and Pielou evenness, ampvis2 (2.7.2) for PCoA ordination, stats (4.1.0) for ANOVA calculations and hierarchical clustering, fpc (2.2-9) for definition of cluster number, factoextra (1.0.7) and ggdendro (0.1.22) for dendrogram creation, DESeq2 (1.32.0) for differential abundance analysis and ggplot2 (2.2.1) and patchwork (1.1.1) for visualization.

### Testing for Behavioral and Physiological Responses

The set of behavioural outcomes belongs to the confirmatory part of the study and appropriate sample size was determined a priori by a power analysis using simulated sampling for a two-way ANOVA design. The power analysis was done for the main outcome variable, plasma corticosterone levels in the SRT. Based on historical data ^13,14^, we expected to observe an effect of medium size (*i.e.* means estimates for two randomly chosen RFs are expected to be in the range of 20%, equivalent to a ratio of between-facility: within-facility variation of 1:2). This resulted in a required minimal sample size of 12 mice per RF and sex. Further information on the sample size calculation is available in the Reporting Summary and Supp. Fig.8. The same sample size (n=12 per RF and sex) was also used for behavioural testing.

The order of behavioural test trials for all mice was randomized using the random number generator of the Mathematica software (version 11; Wolfram Research, Champaign, Illinois, USA). Forty mice (20 males and 20 females) were randomly assigned to each of 3 experimenters. Mice were handled by the same experimenter during the habituation period, cage change and behavioural testing. Animals were tested in parallel (at the same time, but in separate apparatuses) by two experimenters, with 3 different combinations of 2 experimenters each day. Testing was carried out in batches during four consecutive days. Males were tested on the 1^st^ and 3^rd^ day, while females were tested on the 2^nd^ and 4^th^ day. Each day’s testing was done by each experimenter in 3 blocks of 5 animals, each. The randomization and allocation procedures were restricted so that in each block for each experimenter there was exactly one mouse from each RF in random order, with the addition that in no case animals tested at the same time were from the same RF. The allocation and randomization scripts are available as supplementary file.

#### Open field test

The open-field test (OF) was performed in a polycarbonate box (45 x 45 x 45 cm; illumination set to 120 lux) with grey walls and a white base plate. Each mouse was placed in the left (close to the experimenter) corner, facing the wall, and allowed to freely explore the open field for 10 minutes. Recording started immediately after placing the animal in the box. The behaviour of the first five minutes of the test was analysed.

The behaviour was video recorded using an infrared camera system and mice were automatically tracked from videos using EthoVision® XT software (version 11.5; Noldus, Wageningen, Netherlands). The space was virtually divided into a centre zone (20 x 20cm) and an outer zone. The total distance traveled, the average velocity, the number of entries into the center area, the time spent in the center and the latency to the first center entry were scored.

#### Light dark box test

The light dark box test (LDB) was conducted in a box (37.5 x 21.5 x 15 cm) consisting of a small, closed dark compartment (12.5 × 21.5 x 15 cm; illumination set to 5 lux) and a larger light compartment (25 × 21.5 x 15 cm; illumination set to 200 lux) connected by a sliding door. Each mouse was placed in the dark compartment of the apparatus and testing began as the sliding door to the light side of the box was raised, and the duration of the test was 10 minutes. The behaviour of the first five minutes of the test was analysed. The total distance traveled, the average velocity in the light compartment, the time spent in the light compartment, the number of entries into the light compartment and the latency to enter the light compartment were measured from video recordings using EthoVision® XT software (version 11.5; Noldus, Wageningen, Netherlands).

#### Stress reactivity test

The stress reactivity test (SRT) was performed according to established protocols ^15^ with slight modifications. In brief, each mouse was taken out of its home cage and a first blood sample was collected by incision of the ventral tail vessel with a scalpel blade (Paragon® disposable sterile scalpels No. 10, Paragon Medical, Lausanne, Switzerland). The procedure was limited to 2 minutes to obtain basal levels of corticosterone unaffected by the sampling procedure. Immediately after blood collection, the mouse was restrained for 20 min in a 50 ml plastic conical tube (11.5cm x 2.5 cm; Fisherbrand™ Easy Reader, Fisher Scientific AG, Reinach, Switzerland) with custom-made holes for breathing and for the tail. At the end of the 20-min restraint period, a second blood sample was taken from a fresh incision rostral to the first one, followed by placing the mouse back in its home cage. 90 minutes after the onset of restraint, a third blood sample was taken from a third incision rostral to the second one. Each blood sample was collected by using a dipotassium-EDTA capillary blood collection system (Microvette® CB 300 K2E, Sarstedt, Nümbrecht, Germany). Immediately after sampling, the blood samples were placed on ice. Within 60 minutes, the samples were centrifuged for 10 minutes at 4000*g* and 4°C. Plasma samples were transferred to new, labeled microcentrifuge tubes and stored at −80 °C until assayed. Plasma concentrations of corticosterone were determined by a commercial ELISA kit (EIA 4164, DRG Instruments GmbH, Marburg, Germany) in duplicates according to the manufacturers’ instructions. Intra-and inter-assay coefficients of variation were below 10% and 12%, respectively.

### Oestrous cycle determination

The oestrous cycle stage was assessed by cytological analysis of vaginal smears to estimate the sex hormone status of the female mice. Vaginal smears were taken immediately after testing and post-mortem after sacrificing. Briefly, after each test, the female was placed on the cage lid with its hind end towards the experimenter. The rounded tip of a disposable pipette with 50 μl of sterile distilled water was gently placed at the opening of the vaginal canal and vaginal smear cells were collected by lavage. Smears were placed on microscopic slides, allowed to dry, stained with 0.1% crystal violet solution, washed, and then analysed using light microscopy. The stage of the oestrous cycle was determined based on the relative ratio of nucleated epithelial cells, cornified squamous epithelial cells and leukocytes. Since there were uneven distributions of oestrous cycle stages across groups on any given testing day ^16^, the vaginal smears data were combined into high-oestrogen state (proestrus, dioestrus-proestrus transition, and proestrus-oestrus transition) and low-oestrogen state (dioestrus, oestrus, metoestrus, oestrus-metoestrus transition, and metoestrus/dioestrus transition) and were included in the analysis as a linear binary factor.

### Statistical analysis of behavioural and physiological responses

All statistical analyses were performed using the statistical software R (version 3.6.2). A detailed R script is available as a supplementary file. Data of male and female animals were analysed separately. Statistical tests and models employed for each analysis together with information on fixed and random factors are reported in the Extended Data Tables 3-8.

In brief, linear models without interaction terms and with identity link function were run for the body weight data collected at TP1 (*i.e.* right before euthanasia within each RF; n=6 mice per rearing lab and sex; sample size was limited by the minimal number of cages with 3 mice per sex and RF) and for plasma corticosterone levels measured in the SRT (n=12 mice per RF and sex). Linear mixed models without interaction terms and with identity link function were run for body weight data and for relative adrenal weights (n=24 per RF and sex) collected at TP2.

Satterthwaite approximation was used for determination of *p*-values in the mixed models. A Dunn-Sidak Bonferroni correction method was applied to correct for multiple testing where necessary. For the physiological measures, such as body weight and plasma corticosterone levels, the threshold was set to α’=1-(1-0.05)^1/3^=0.0169.

The distribution of the observed values for behavioural outcomes was inspected for deviations from normality with Q-Q plots. The dataset of the physiological measures (body weights, relative adrenal weights, and corticosterone responses in the SRT) were normally distributed (Suppl. Figs. 9-11). Due to skewed distributions, transformations of behavioral data were necessary for seven variables (Suppl. Fig. 12). In the male cohort, OF latency, LDB latency and LDB time in the light of males were square-root transformed (Suppl. Fig. 12a). In the female cohort, OF distance was log transformed, OF time in the center, OF latency and LDB time in the light were square-root transformed, while LDB latency was arcsine transformed (Suppl. Fig. 12b).

For the analysis of behavioral outcomes, we used a multivariate analysis of variance (MANOVA) without nesting and interaction terms and with identity link function. To check for correlation between recorded variables, we calculated Pearson’s product moment correlation coefficient. We first excluded variables that were highly correlated (Suppl. Table 2), which resulted in the final list of six dependent behavioral variables (OF distance, OF time center, OF latency, LDB time light, LDB entries into light, LDB latency). Pillai’s Trace was used to evaluate the MANOVA differences, while the robustness of the findings was confirmed by three other test statistics: the Wilk’s Lambda, Hotelling-Lawley Trace, and Roy’s Largest Root.

Multivariate outliers were identified by using the squared Mahalanobis distance (mvoutlier.CoDA version 2.0.9; ^17^). The analysis suggested the existence of 6 outliers in the male cohort and 6 in the female cohort; Suppl. Fig. 13), which were not removed for the further analysis. To investigate whether the outliers had an impact on the results, we re-run the analysis with the outliers removed and could confirm that results were not markedly different.

### ATAC-seq analysis on purified neuronal nuclei

#### Nuclei isolation and fluorescence-activated nuclei sorting (FANS)

The ATAC-seq analysis was performed on ventral hippocampi isolated from male mice at TP1 (PND 56) and TP2 (PND 102). For this, test-naïve male mice were used to avoid effects of testing on the chromatin profile. Five cages per RF were selected randomly from all cages containing 3 male littermates after weaning. One mouse from the selected cages was sacrificed in the RF at TP1 (total n=25, i.e. 5 biological replicates per rearing laboratory), while its test-naïve littermate was sacrificed in the testing laboratory at the end of the experiment (TP2; total n=25, i.e. 5 biological replicate per RF).

The analysis has been restricted to males because they showed the most pronounced phenotypic differences in behaviour. Furthermore, in the female cohort we would not be able to distinguish between the differences in chromatin organization induced by hormonal fluctuations ^18^ and differences induced by differential rearing environments.

Total nuclei isolation and purification of neuronal nuclei were performed as described elsewhere ^18,19^ with slight modifications. In brief, the ventral hippocampus was dissected from one side of the brain at −20C°, cut into small pieces and stored in Eppendorf DNA LoBind 2mL tubes at −80°C until nuclei preparation. Nuclei preparation and sorting was performed in 5 batches per each time point, with each batch having exactly one samples from each facility in random order.

To obtain fresh nuclei, frozen tissue samples were resuspended in 4 ml of tissue lysis buffer (0.32 M Sucrose, 5 mM CaCl_2_, 3 mM Mg(CH3COO)_2_, 0.1 mM EDTA, 10 mM TRIS-HCl (pH 8), 1 mM DTT, 0.1% Triton X-100) and dissociated by 30 strokes of pestle A (loose pestle) and then 20 strokes of pestle B (tight pestle) in a glass douncer (7 ml Dounce tissue grinder set, KIMBLE, DWK Life Sciences). The lysate was transferred to an ultracentrifuge tube, followed by adding 6 ml of sucrose buffer (1.8 M Sucrose, 3 mM Mg(CH_3_COO)_2_, 1 mM DTT, 10 mM TRIS-HCl pH 8) underlaid beneath the solution. The samples were then spun at 171,192.8×g in a Hitachi Ultracentrifuge (CP100NX; with Sorvall TH-641 swing bucket rotor) for 1 hour at 4°C. Next, the nuclei pellet was resuspended with 500μl of 0.1% BSA in DPBS with glucose, sodium pyruvate, calcium, and magnesium. The nuclei solution was than incubated with monoclonal antibody against neuronal marker NeuN conjugated to AlexaFluor®488 (1:1000; Merk Millipore, MAB377X) for one hour at 4°C on rotation protected from light. After incubation, DAPI (1:1000; ThermoFisher Scientific, 62248) was added to the reaction. The nuclei suspension was immediately taken to be processed on a FACSAria instrument (BD Biosciences, USA) at the Flow Cytometry and Cell Sorting Core Facility (FCCS CF) of the Department for BioMedical Research, University of Bern.

Prior to sorting, samples were filtered through a 35 μm cell strainer. To set up the gating protocol we used four controls: i) unstained nuclei only ii) DAPI only; iii) IgG1 isotype control-AlexaFluor 488 and DAPI; and iv) NeuN-AlexaFluor 488 only; in addition to a sample containing NeuN-AlexaFluor 488 and DAPI stain (Suppl. Fig. 14), which allowed us to eliminate debris and any clumped nuclei effectively, resulting in an apparent separation of the NeuN + (neuronal) nuclei populations. From each individual ventral hippocampus, we collected 80,000 NeuN+ (neuronal) nuclei in BSA-precoated tubes filled with 200 μL of DPBS. The purity of the sorted nuclei was confirmed by re-sorting a small fraction of NeuN+ nuclei using the same protocol (showed more than 98% purity).

#### Transposition reaction, ATAC-seq libraries preparation and sequencing

ATAC-seq was performed according to Buenrostro and colleagues^20^, with some modifications. Following FANS, neuronal nuclei from the ventral hippocampus were spun down (2900×g, 10 minutes at 4°C). The supernatant was carefully removed, avoiding the visible nuclei pellet. The nuclei pellet was resuspended in 50 μl of the transposase reaction mix including 25 μL 2×TD reaction buffer and 3 μl Tn5 Transposase, (Illumina Tagment DNA TDE1 Enzyme and Buffer Kits, 2003419) and 22 μl of nuclease free water (NFW; Ambion™, AM9937). The transposition reaction was performed at 37°C for 30 minutes in a thermomixer with 500 RPM mixing, followed by purification using a MinElute PCR Purification Kit (Qiagen, 28004). The purified, transposed DNA was eluted in 10 *μl* of EB elution buffer and stored at *-20°C* until amplification.

To amplify transposed DNA fragments, the following procedure was performed in two batches of 25 samples (one batch per each time point) with equal group distribution (n=5 samples from each RF). For indexing and amplification of transposed DNA, we combined the following for each sample: 10 μl transposed DNA, 25 μl NEBNext High-Fidelity 2× PCR Master Mix (New England Biolabs, M0541S), 9 μl of unique, dual-indexed primer (IDT for Illumina Nextera DNA UD Indexes; 20026930) and 6 μl of NFW (Ambion™, AM9937). The PCR reaction was carried out using the following conditions: 1 cycle of 72°C for 5 minutes and 98°C for 30 seconds; followed by 5 cycles of 98°C for 10 seconds, 63°C for 30 seconds, and 72°C for 1 minute; and a hold step at 4°C.

We then performed a qPCR side reaction to manually assess the amplification profiles and determine the required number of additional PCR cycles ^21^. The reaction mix was prepared by combining 5 μL of a previously PCR-amplified DNA with, 7.5 μl of SYBR® Green PCR Master Mix (Applied Biosystems, 4344463) and 2.5 μl of NFW, and cycling conditions were set as follows: 1 cycle of 98°C for 30 seconds, followed by 20 cycles of 98°C for 10 seconds, 63°C for 30 seconds, and 72°C for 1 minute. Under our experimental conditions, 2-4 PCR cycles were added to the initial set of 5 cycles. The amplified libraries were purified using MinElute PCR Purification Kit (Qiagen, 28004) and eluted in *20* μL of the EB elution buffer. Library quality was monitored using the Advanced Analytics Fragment Analyzer CE12 (Agilent, USA; Suppl. Fig. 15), and the concentration was determined by Qubit HS DNA kit (Life Technologies, Q32851) and quantitative PCR with the library quantification kit from Bioline Jet Set Library Quantification Kit LoROX (Meridian Bioscience, BIO-68029).

A total of 50 ATAC-seq libraries was sequenced in two batches with equal group distribution (25 libraries/batch/ NovaSeq S1 Flow Cell; n=5 per each rearing laboratory) on the Illumina NovaSeq 6000 instrument with 2×100 bp pair-end protocol at the Next Generation Sequencing (NGS) Core Platform of the University of Bern.

#### ATAC-seq Data Analysis

Sequencing reads were trimmed, and quality checked with fastp (version 0.20.1 ^22^) with CTGTCTCTTATACACATCT as adapter sequence and a minimal read length of 30 bp. Reads were aligned to the mouse reference genome (ensembl build 102) with Bowtie2 in paired end mode (version 2.3.5.1; ^23^) keeping only concordant and unique alignments.

Duplicate read pairs were marked using the *MarkDuplicates* command from the Picard software suite (version 1.140; broadinstitute.github.io/picard/). Peaks were then called in each sample separately with MACS2 (version 2.1.4, with the parameters *-f BAMPE -g mm --nomodel -q 0.05 --broad --broad-cutoff 0.1 --keep-dup all)* as previously reported ^24^. For each time point and RF, peaks were intersected with multovl (version 1.3; ^25^ and only peaks found in at least three samples per group (*i.e.* rearing laboratory) were kept (Suppl. Fig 16). Finally, peaks from all groups were merged with multovl (union of all peaks within groups). The number of reads within peak intervals was obtained with featureCounts (version 2.0.1;^26^) with the parameters *--primary --ignoreDup --minOverlap 30.* The peak set was further used to build a distance matrix using the aligned reads of the individual samples per each time point. The sample correlation matrices were generated for by using all sites and 10% of the most variable sites and visualized by correlation heatmaps, PCA and t-SNE distance plots.

The distance matrix was also used as an input for the PERMANOVA. By using PERMANOVA (function adonis() from the R-package vegan version 2.5-7 ^27^, we tested whether and to what extent the variation between samples can be explained by the RF within each time point. Since the test was based on 9999 permutations, the lowest possible *p*-value was set to “< 0.0001” instead of “0”.

Variation in read counts was analyzed with a general linear model in R (version 3.6.1) with the package DESeq2 (version 1.24.0; ^28^) according to a factorial design with the two explanatory factors “RF” and “processing batch”, within each timepoint. For the annotation of peaks, we used the ChiPseeker annotation for the plot with genomic features and the Homer annotation for TSS distance and candidate, protein coding, genes. Following specific conditions were compared with linear contrasts: i) one-to-one (oto) comparison of each pair of laboratories (RF1 vs RF2, RF1 vs RF3, etc) for each time point; ii) one-to-many (otm) comparison of one laboratory to all other laboratories for each time point (RF1 vs all other RFs, RF2 vs all other RFs, etc), and iii) a global test for the factor “RF” (LRT_RF), *i.e.* do different rearing laboratories differ in general.

Within each comparison, *p-values* were adjusted for multiple testing (Benjamini-Hochberg), and regions with an adjusted *p-value* (false discovery rate, FDR) below 0.01 and a minimal log2 foldchange (*i.e.* the difference between the log2 transformed, normalized sequence counts) of 0.5 were considered to be differentially accessible. Normalized sequence counts were calculated accordingly with DESeq2 and log2 (x+1) transformed. Gene Ontology (GO) and KEGG (Kyoto Encyclopedia of Genes and Genomes) Pathway analyses were performed on the significant peaks located 2 kb up- and downstream of the transcriptional start site. Functional annotation for enrichment of GO terms was performed using topGO (version 2.28 ^29^) in conjunction with the GO annotation from Ensembl available through biomaRt ^30^. Analysis was based on gene counts using the “weight” algorithm with Fisher’s exact test (both implemented in topGO). Only GO terms with more than 5 genes were tested and terms were identified as significant if the *p*-value was below 0.05. Enrichment of KEGG pathways in gene sets was tested with clusterProfiler (version 3.12.0 ^31^) using the gene to pathway mappings available through biomaRt ^30^ and the package org.Rn.eg.db (version 3.8.2 ^32^). Integrative Genome Viewer (IGV, version 2.8.9) was used to visualize and extract representative ATAC-seq tracks.

## Supplementary Information

### I. Supplementary Figures

**Supplementary figure 1:** Timeline of the study.

**Supplementary figure 2:** Diet suppliers: Differentially abundant taxa.

**Supplementary figure 3**: Oestrous cycle-dependent effects on behavioral phenotype in female mice from different rearing facilities.

**Supplementary figure 4:** Oestrous cycle-dependent effects on basal corticosterone levels in female mice from different rearing facilities

**Supplementary figure 5:** Neuronal chromatin accessibility differs in males from different rearing facilities.

**Supplementary figure 6:** Random down-sampling of 16S sequencing reads.

**Supplementary figure 7:** Evaluation of differences in microbiome between samples from behaviorally tested-BT mice and mice used for chromatin profiling-MA mice.

**Supplementary figure 8:** Power analysis

**Supplementary figure 9:** Q-Q (quantile-quantile) a probability plots for the body weight data points.

**Supplementary figure 10:** Q-Q (quantile-quantile) a probability plots for the corticosterone response data points in the stress reactivity tests.

**Supplementary figure 11:** Q-Q (quantile-quantile) a probability plots for the relative adrenal gland weight at TP2.

**Supplementary figure 12**: Q-Q (quantile-quantile) a probability plots for the behavioural data sets.

**Supplementary figure 13:** Multivariate outliers.

**Supplementary figure 14:** Gating strategy for separation of neuronal nuclei using fluorescence-activated nuclei sorting (FANS).

**Supplementary figure 15:** ATAC-seq library quality control.

**Supplementary figure 16:** ATAC peak count statistics.

### II. Supplementary Tables

**Supplementary Table 1** (given as an Excel file): Detailed housing and husbandry conditions in each rearing facility (a) and testing laboratory (b).

**Supplementary Table 2:** Pearson product moment correlations for behavioural measures for males **(a)** and females **(b)**.

### III. Code Availability (given as a separate PDF file)

**R Markdown 1:** Microbiome

**R Markdown 2:** Behavioural and physiological outcomes

**R Markdown 3:** Chi-squared test (Behavioural Classification)

**Matematica Code**: Test allocation

### IV. Row Data Availability (given as a separate Excel files)

- Jaric et al. B1_Microbiome Metadata
- Jaric et al. B1_FEMALES_BW _TP1
- Jaric et al. B1_MALES_BW_TP1
- Jaric et al. B1_FEMALES_Summary (behavior, CORT, BW only BT mice)
- Jaric et al. B1_MALES_Summary (behavior, CORT, BW only BT mice)
- Jaric et al. B1_MALES_BW and adrenals (all mice) TP2
- Jaric et al. B1_FEMALES_BW and adrenals (all mice) TP2

**Supplementary figure 1.**
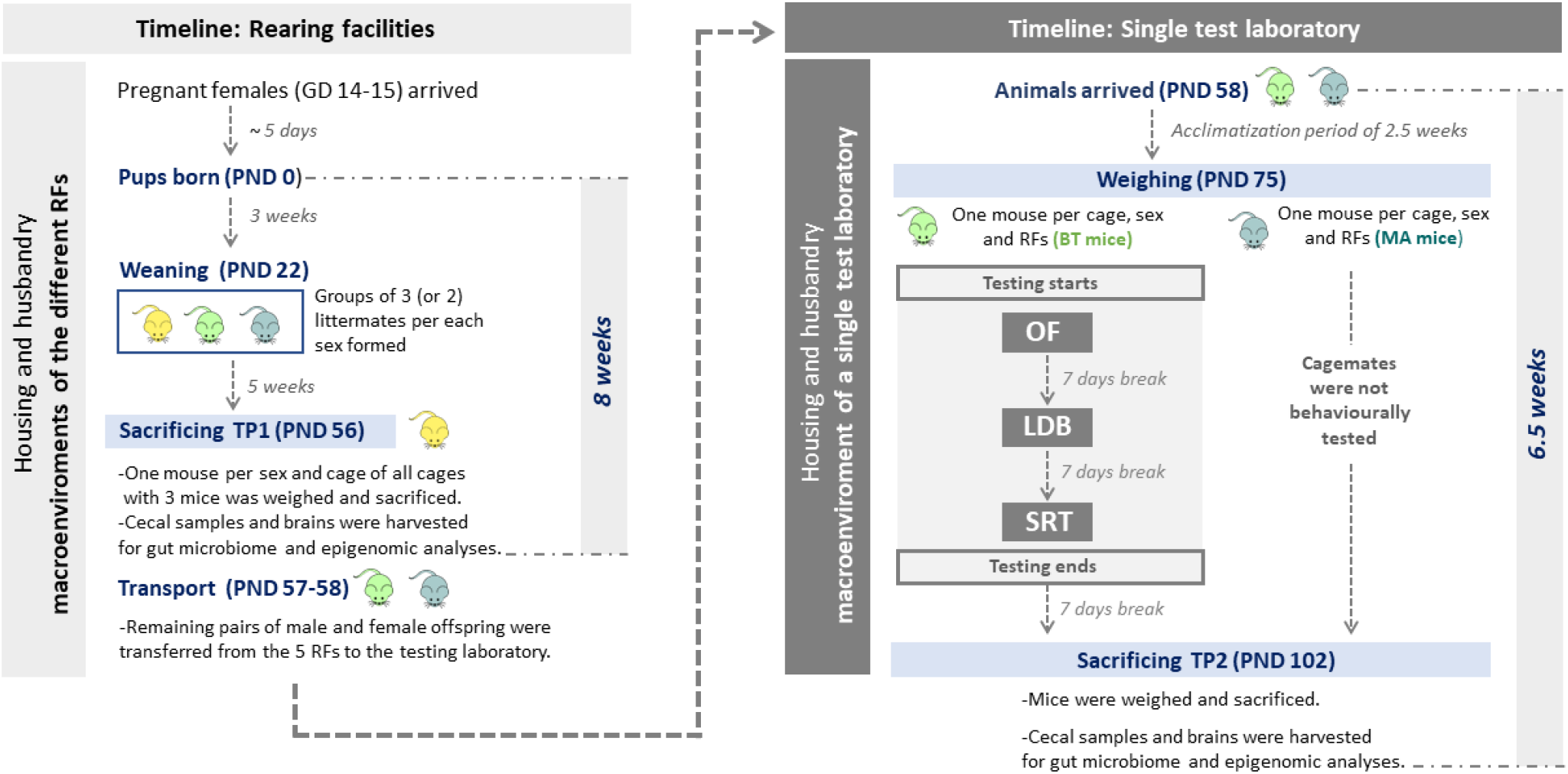
Timeline of the study. GD: gestational day: PND: postnatal day; BT mice: mice used for behavioural testing mice; MA mice: mice that were not behaviourally tested and used for chromatin profiling only. OF: open field test; LDB: light-dark box test; SRT: Stress reactivity test.

**Supplementary figure 2.**
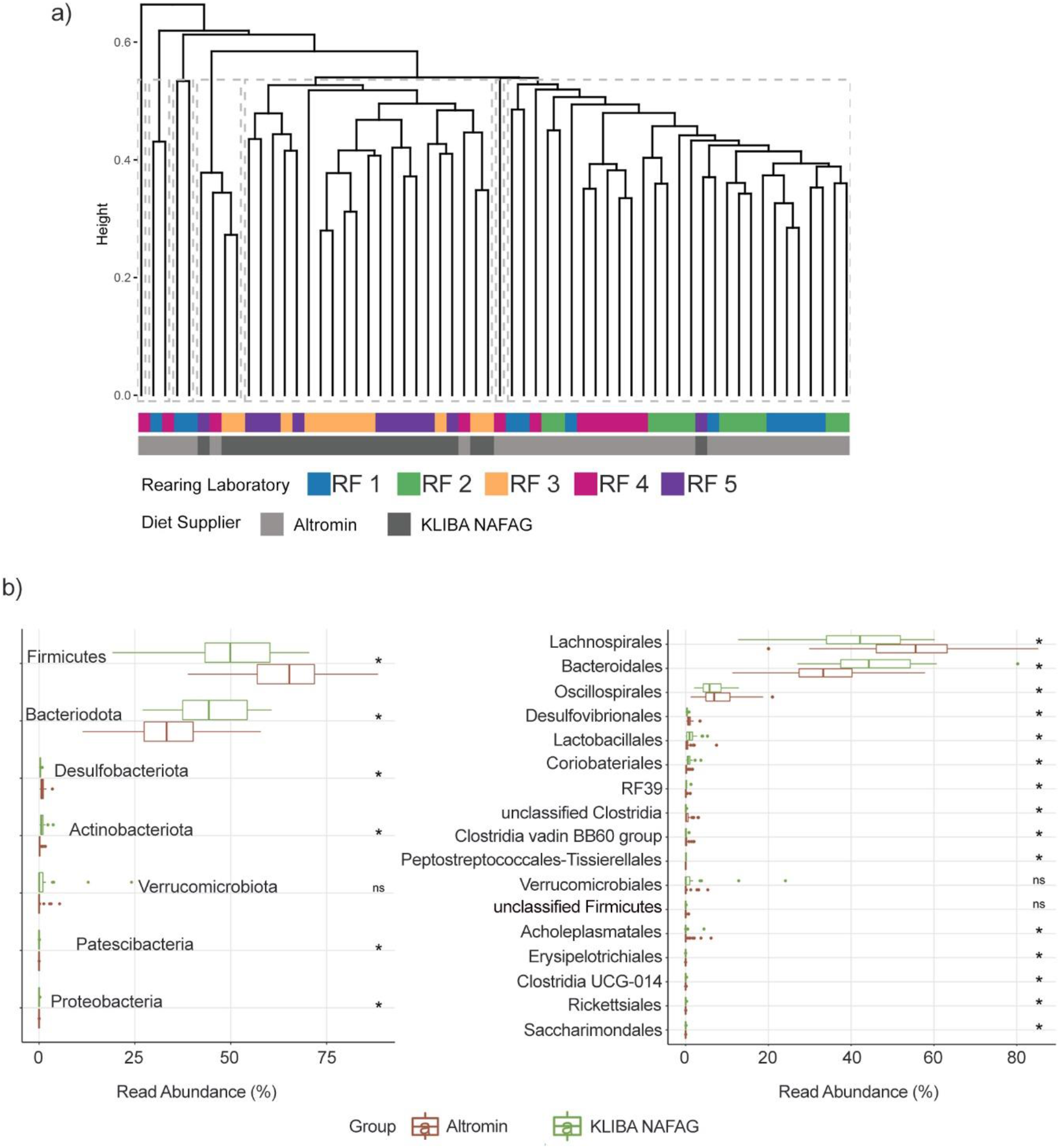
Diet suppliers: Differentially abundant taxa. **a)** Dendrogram representing relationship between samples from mice at TP1 following hierarchical clustering (average linkage). Horizontal bars below the dendrogram represent sample identity in relation to rearing facility (first bar) or diet supplier (second bar). **b)** Mean abundance of amplicon sequence variants (ASVs) identified as differentially abundant between the two diet suppliers aggregated by phylum (left) and order (right). Significance symbols represent results from Wilcoxon paired-rank test.

**Supplementary Figure 3.**
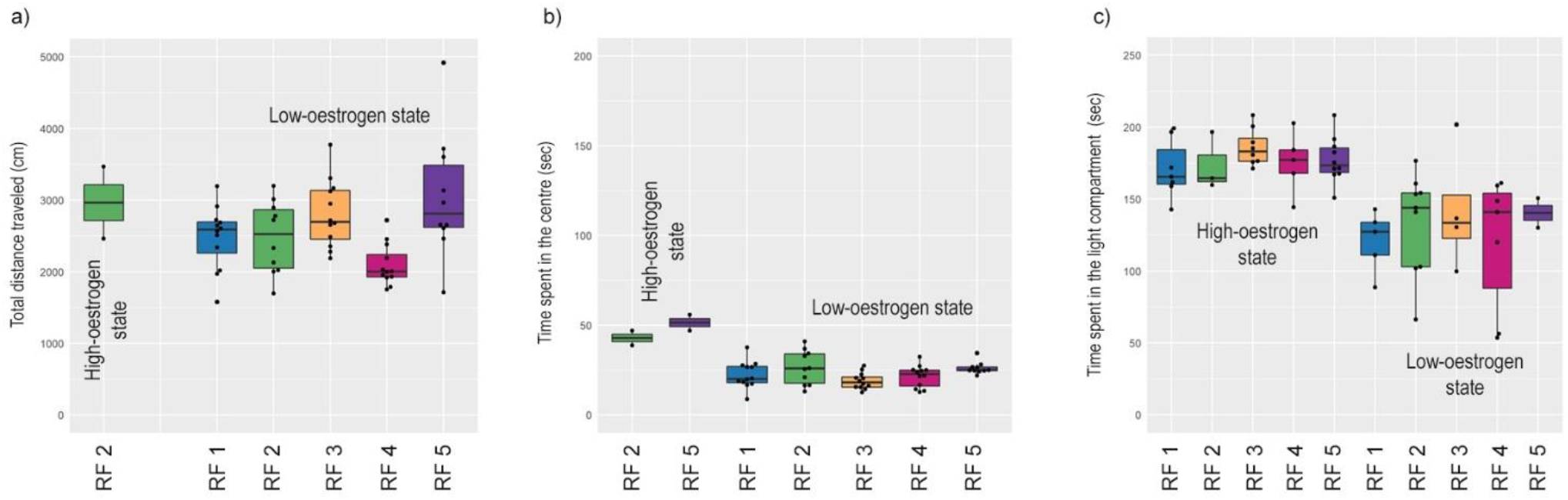
Oestrous cycle-dependent effects on behavioral phenotype in female mice from different rearing facilities. Results of the open field **(a, b)** and light-dark box **(c)** tests are presented in females depending on the oestrous cycle stage determined immediately after behavioral testing. There was a significant effect of oestrogen status on time spent in the center (b) and time spent in the light compartment **(c)** with high-oestrogenic females showing marginally higher activity than low-oestrogenic females.

**Supplementary Figure 4.**
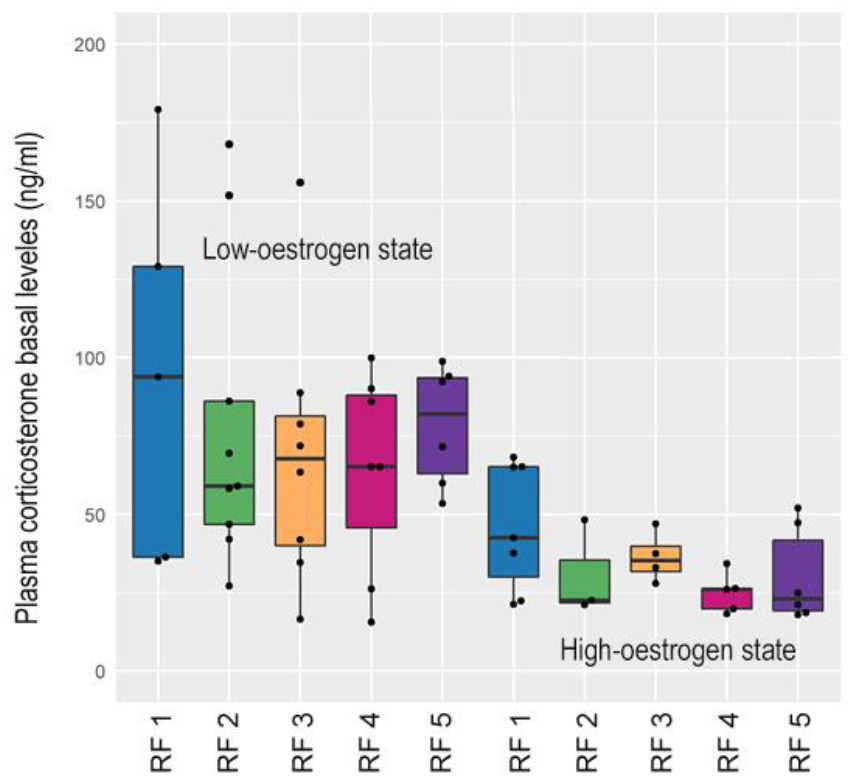
Oestrous cycle-dependent effects on basal corticosterone levels in female mice from different rearing facilities. Results are presented depending on the oeestrous cycle stage determined after HPA reactivity tests. There was a significant effect of oestrogen status on basal corticosterone levels, with high-oestrogenic females showing lower basal corticosterone levels than low estrogenic females.

**Supplementary figure 5.**
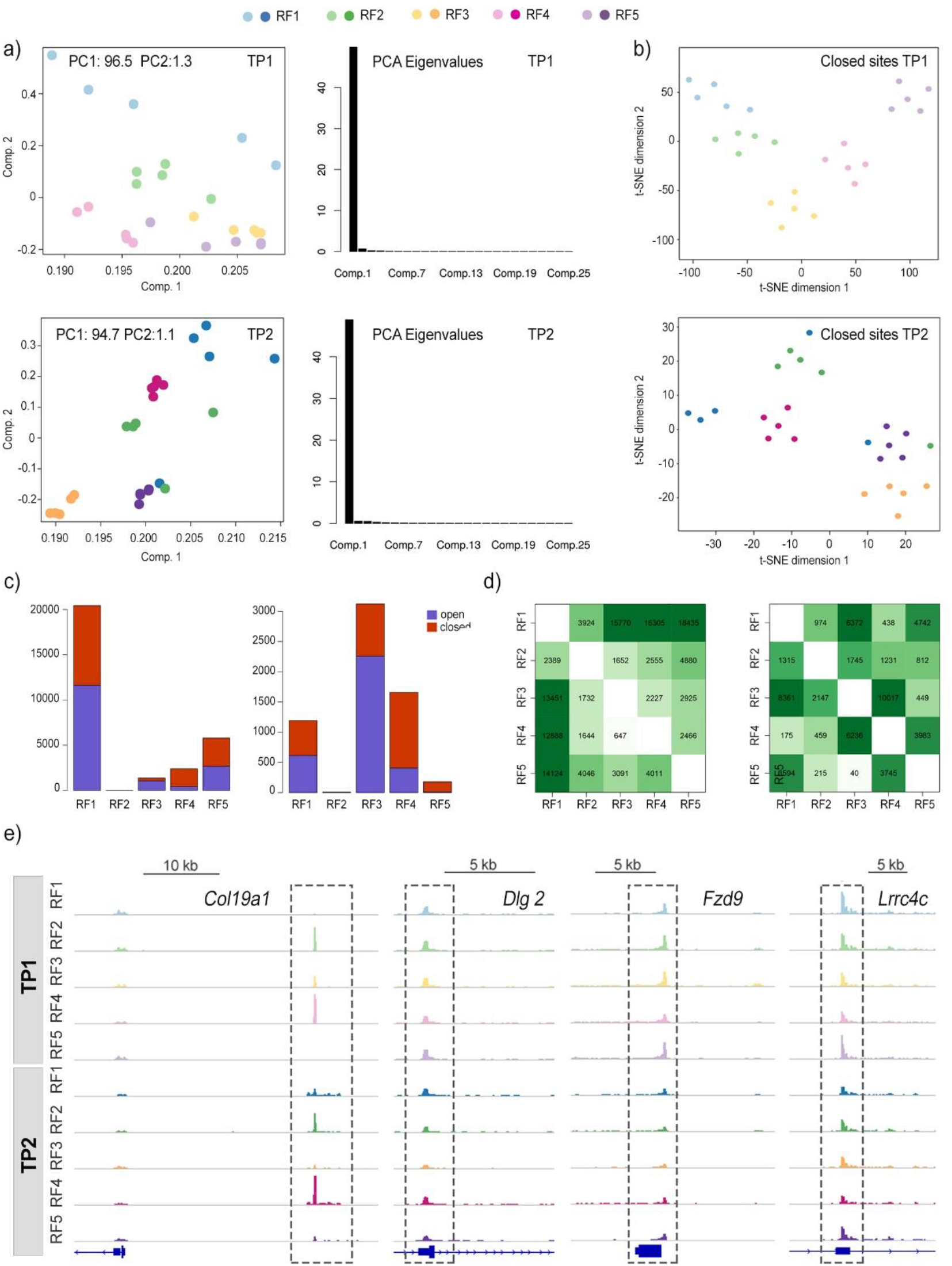
Neuronal chromatin accessibility differs in males from different rearing facilities. **a)** Principal component analysis of the ATAC-seq data. **b)** Manhatten distances between samples for closed chromatin sites visualized by t-SNE. **c)** The number of significant differentially accessible peaks associated to each of rearing environments in a given timepoint (one-to-many comparison; adjusted for multiple testing FDR < 0.01 and abs(logFC) > 0.5. **d)** The number of significant differentially accessible peaks between groups in a given timepoint (one-to-one comparison; adjusted for multiple testing FDR < 0.01 and abs(logFC) > 0.5.) **e)** Chromatin accessibility profiles of the *Col19a1, Dlg 2, Fzd9* and *Lrrc4c.* Shown are genomic coordinates of differential ATAC-seq peaks.

**Supplementary figure 6.**
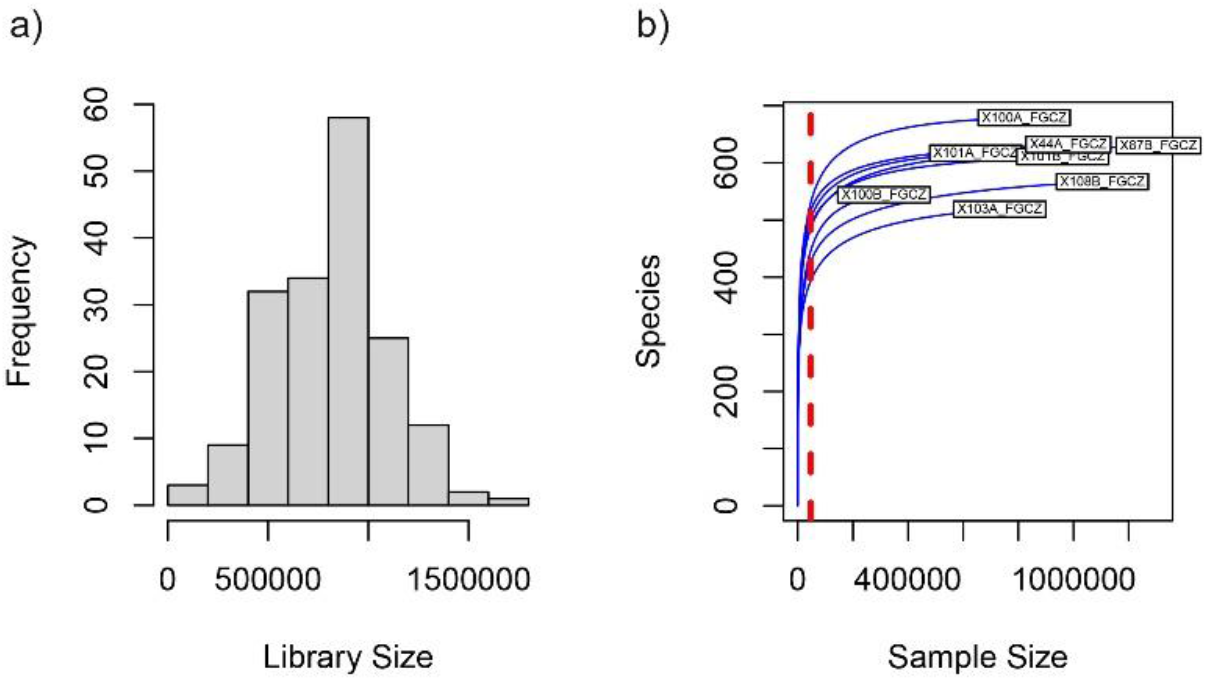
Random down-sampling of 16S sequencing reads. **a)** Histogram of individual library sizes. **b)** Example rarefaction plots of 8 randomly selected samples. The red line indicates rarefaction depth.

**Supplementary figure 7.**
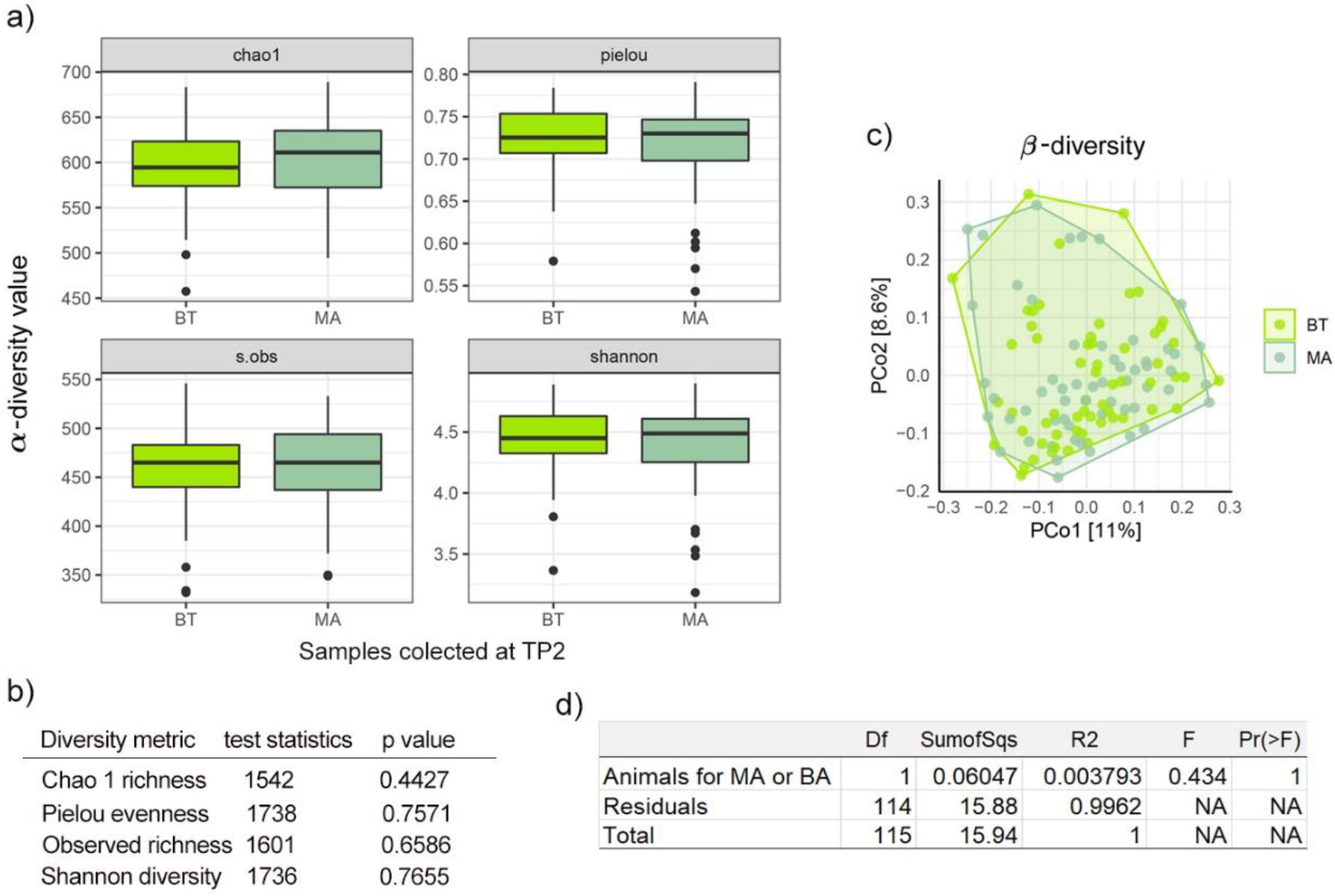
Evaluation of differences in microbiome between samples from behaviorally tested-BT mice and mice used for chromatin profiling-MA mice. **a)** Mean values for α-diversity metrics for MA-mice and BT-mice. Top left-Chao1 richness, bottom left-Observed species richness, top right-Pielou evenness, bottom right-Shannon diversity index. **b)** Results of Wilcoxon signed-rank test for each α-diversity metric between MA and BT mice. **c)** Ordination plot visualizing Principal Coordinate analysis based on Bray-Curtis dissimilarity between samples from MA and BT-mice collected at TP2. **d)** Result of PERMANOVA partitioning variation in microbiome composition between mice used for MA and BT.

**Supplementary Figure 8.**
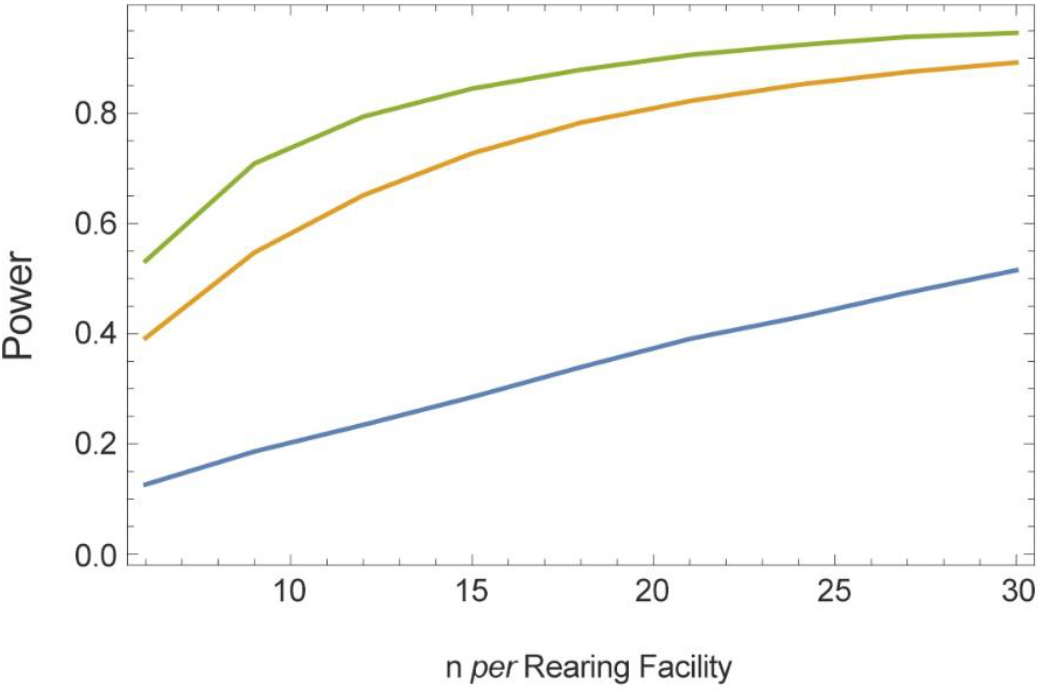
Power analysis: Power curve for a one-way ANOVA with α=0.05, k=5 rearing facilities and n=6 to 30 subjects, assuming an average means difference between two randomly chosen labs of 10% (blue), 20% (orange) or 30% (green). Power estimates are based on 10, 000 repeated samples. To generate differences between labs we sampled distribution means from a normal distribution with the reported mean (10,000 square units) and standard deviation of 890, 1780, and 2260 square units, which resulted in samples, where the difference in the effect size between two randomly chosen labs was on average 10, 20, or 30 percent.

**Supplementary Figure 9.**
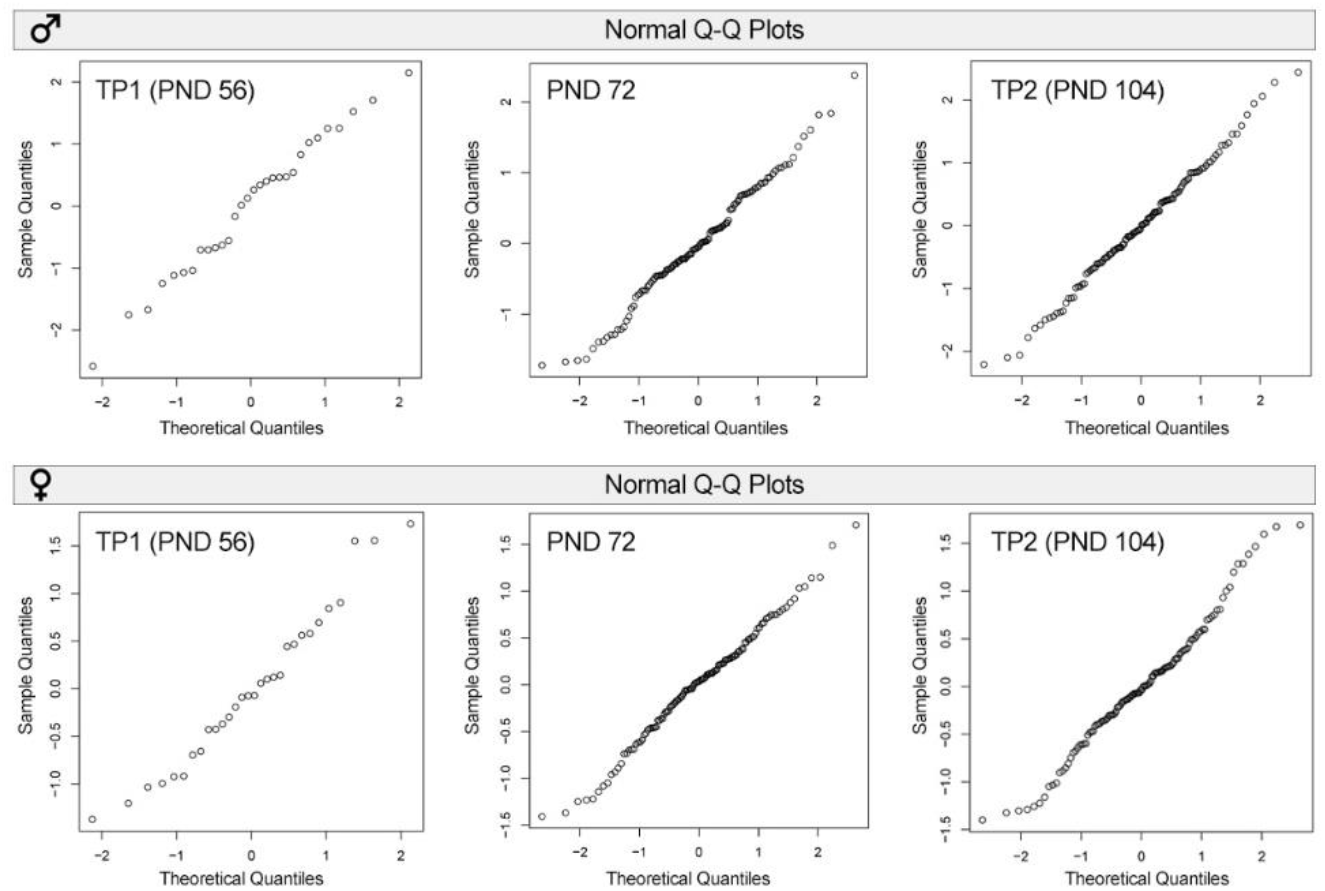
Q-Q (quantile-quantile) a probability plots for the body weight data points. The data points of the body weights were normally distributed both in males (top panel) and females (bottom panel).

**Supplementary Figure 10.**
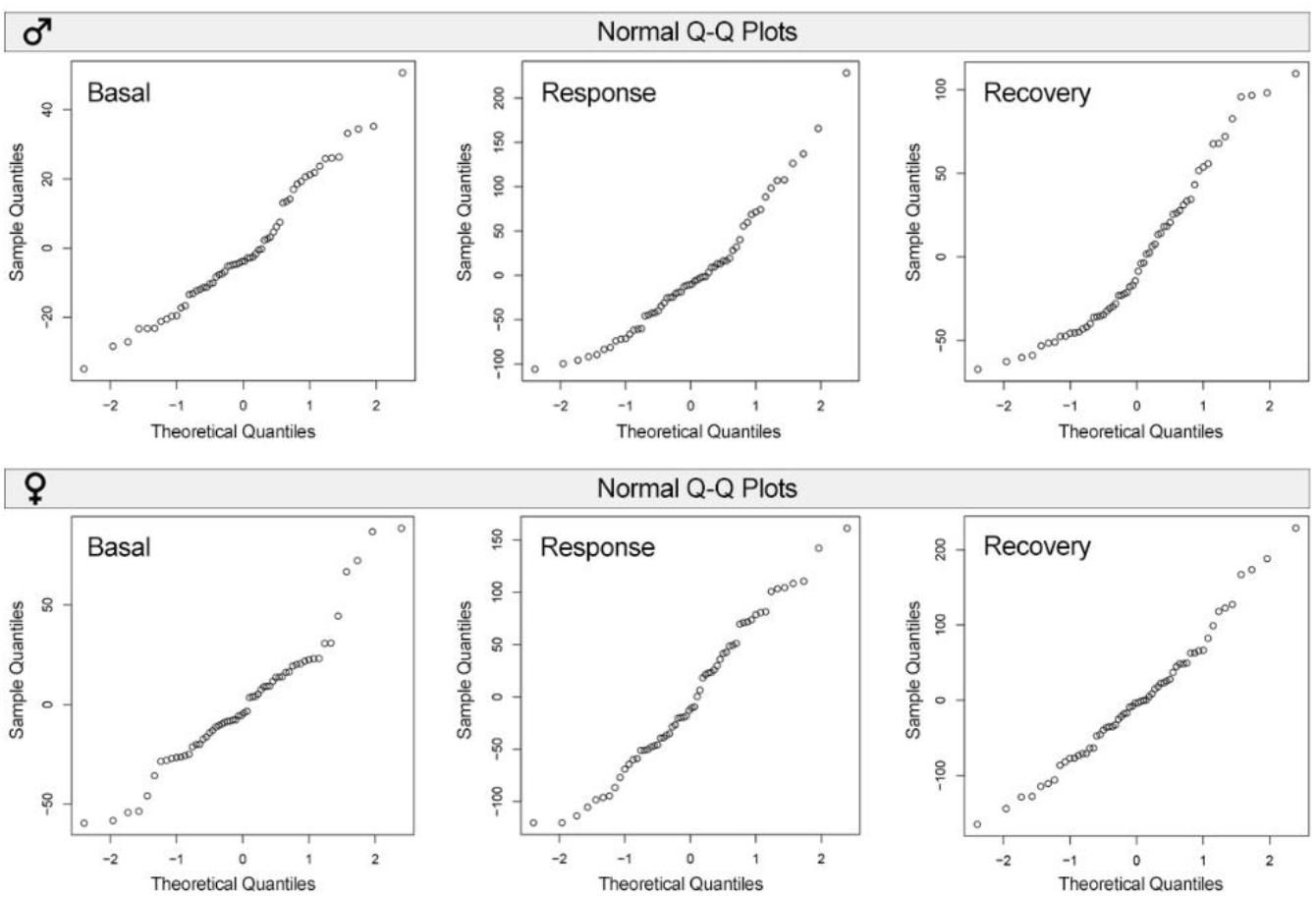
Q-Q (quantile-quantile) a probability plots for the corticosterone response data points in the stress reactivity tests. The data points of the corticosterone response were normally distributed both in males (top panel) and females (bottom panel).

**Supplementary Figure 11.**
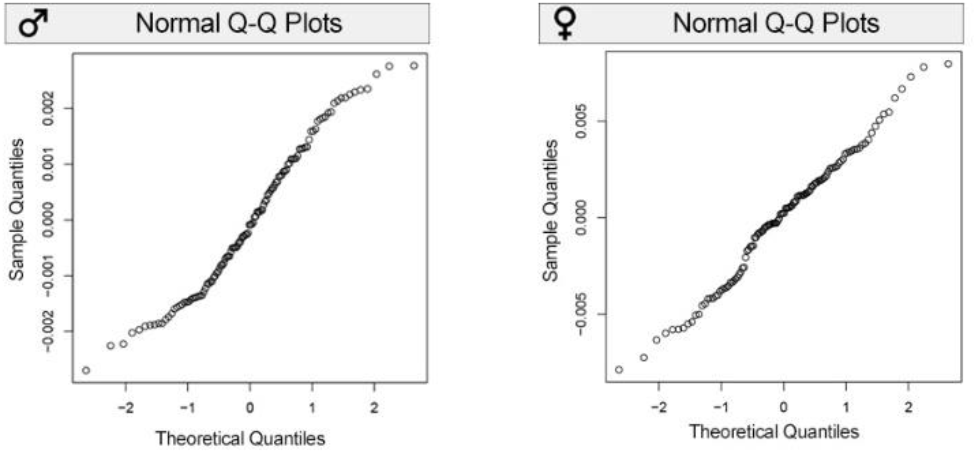
Q-Q (quantile-quantile) a probability plots for the relative adrenal gland weight at TP2. The data points of the relative adrenal gland weights were normally distributed both in males (left plot) and females (right plot).

**Supplementary Figure 12.**
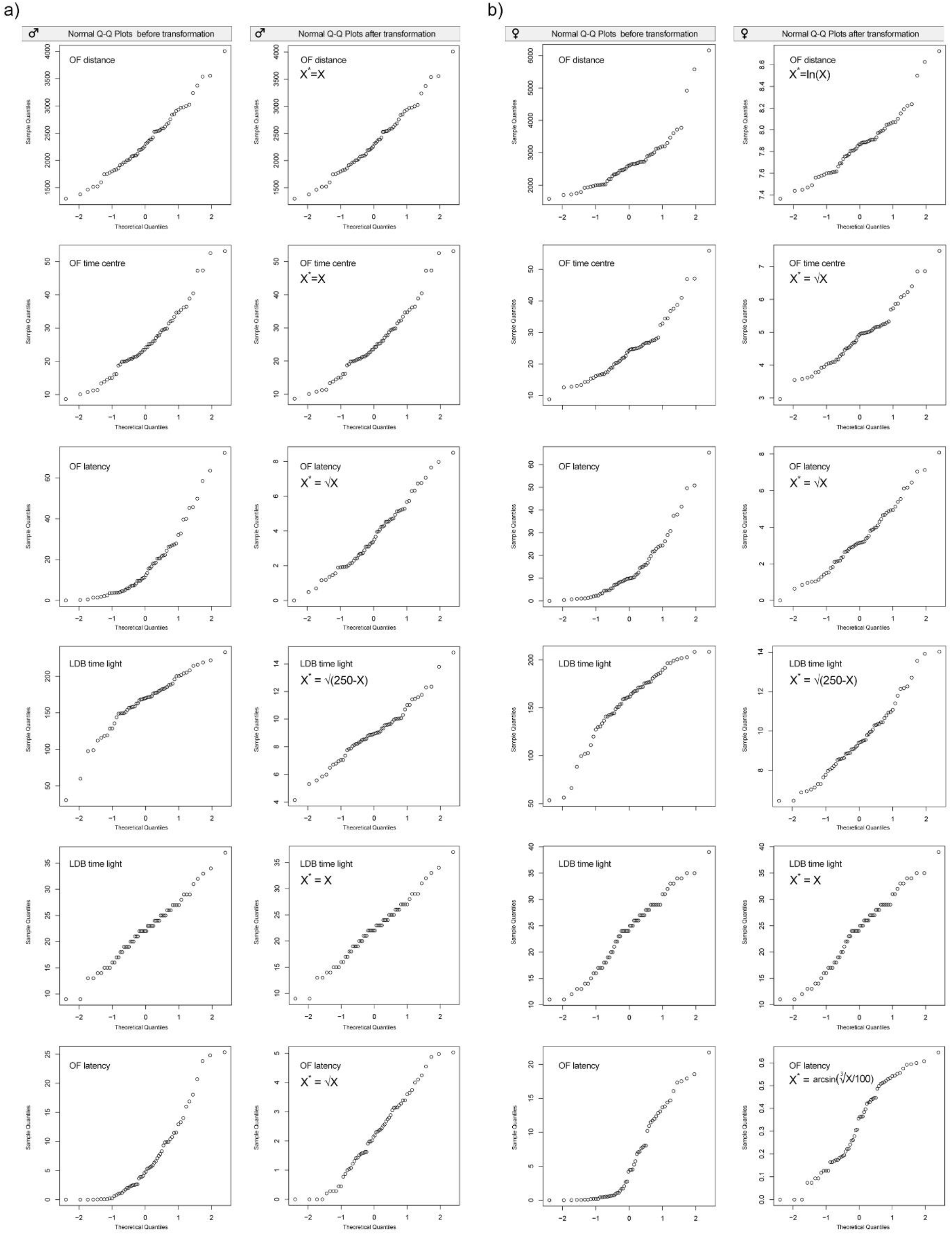
Q-Q (quantile-quantile) a probability plots for the behavioral data sets. Q-Q plots are presented before and after transformation of individual data points for both males **(a)** and **(b)** females.

**Supplementary Figure 13.**
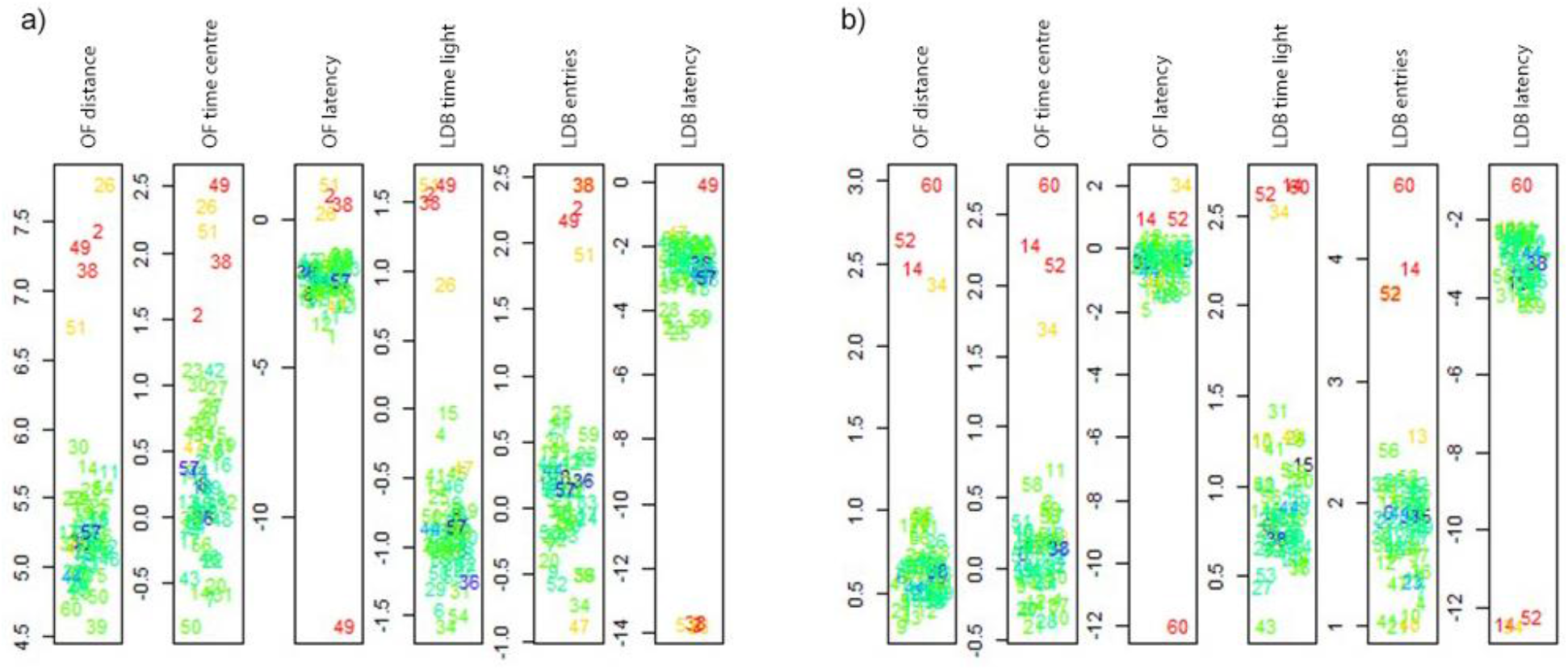
Multivariate outliers. Results of the test for multivariate outliers indicated the existence of 3 outliers in males **(a)** and 3 outliers in the female data **(b)**. Outliers are shown in red.

**Supplementary Figure 14.**
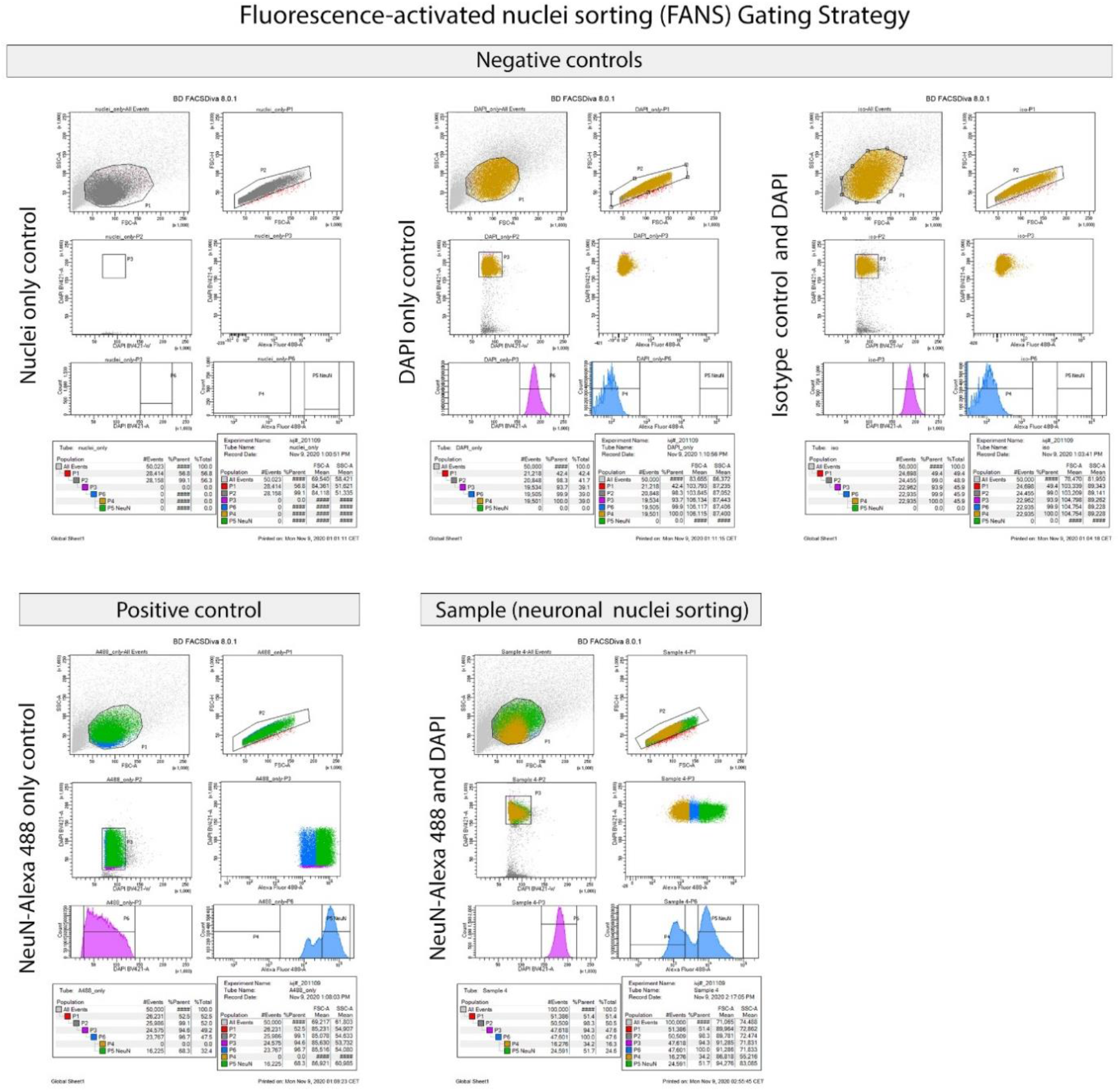
Gating strategy for separation of neuronal nuclei using fluorescence-activated nuclei sorting (FANS). Sorting plots from **three negative controls** (Nuclei only, DAPI only and Isotype control + DAPI) processed without primary antibody (neuron-specific marker NeuN), **positive control** containing NeuN antibody conjugated with Alexa 488 only (NeuN-Alexa 488 only control) and our **sample** processed with NeuN-Alexa 488 antibody and DAPI are shown. Representative FANS reports showing the gating strategy for the checking the size and granularity, removal of debris and ensuring a successful separation of NeuN+ (neuronal) from non-neuronal single nuclei.

**Supplementary Figure 15.**
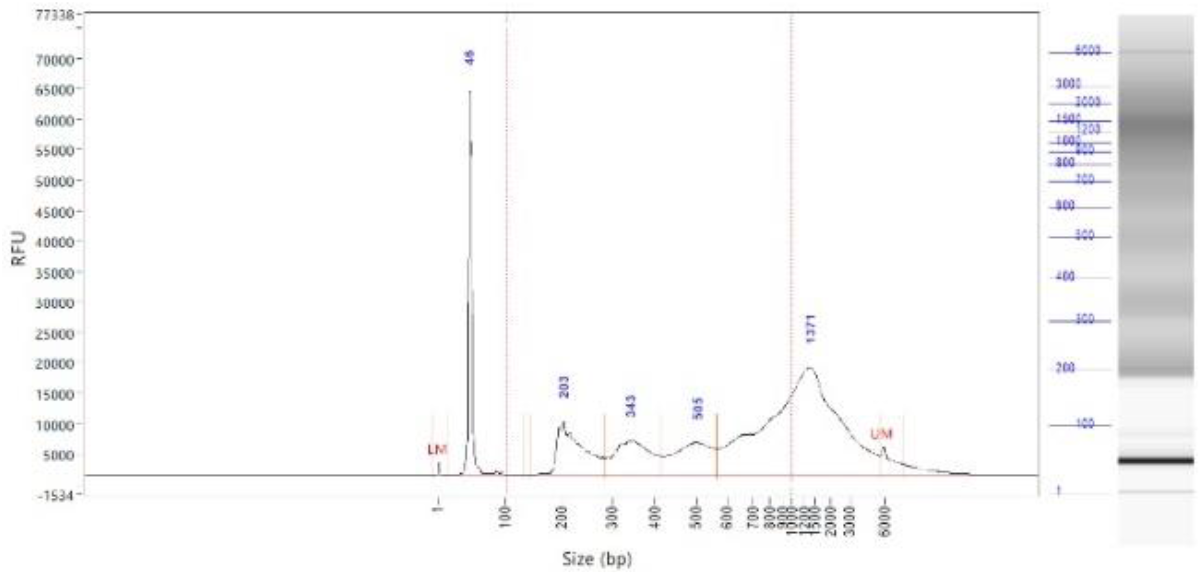
ATAC-seq library quality control. The quality control of ATAC-seq libraries was performed Fragment Analyzer (FA). The representative FA trace with nucleosomal banding pattern is shown.

**Supplementary Figure 16.**
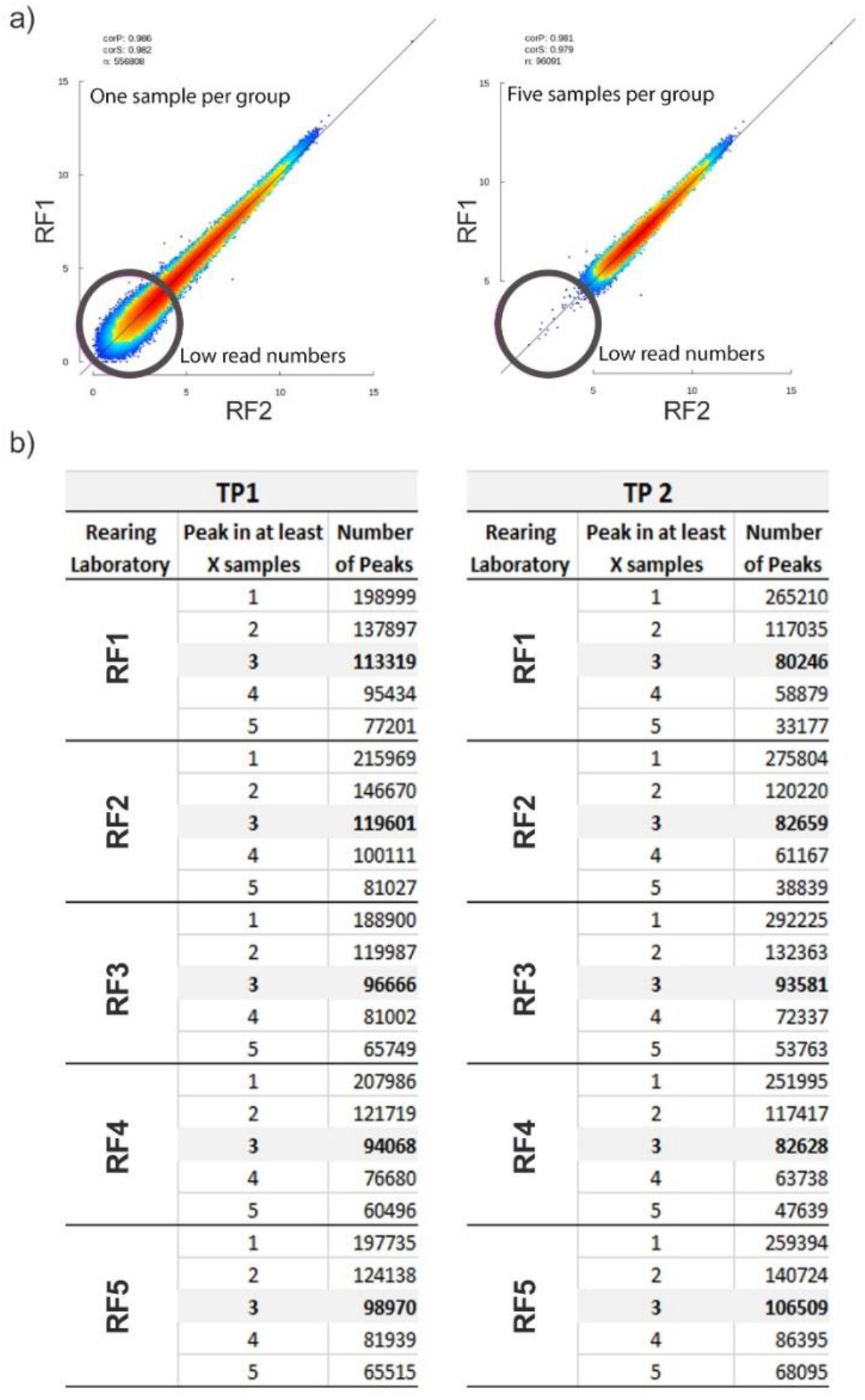
ATAC peak count statistics. The number of peaks drops quite strongly with the minimal number of samples required for merging peaks. a) Representative scatter plots showing log2 (x+1) transformed, normalized values averaged in RF1 and RF2 at TP1. b) Only high-confidence broad peaks, shared across at least three biological replicates of one group are used for all downstream analyses.

**Supplementary Table 2:** Pearson product moment correlations for behavioural measures for males **(a)** and females **(b)**. Several highly correlated variables were removed before a multivariate analysis. Only variables with moderated correlations amongst each other were used. **a)** Pearson product moment correlations for behavioural measures for males. Mean absolute correlation coefficient for males was 0.20.

| MALES | OF distance | OF velocity | OF time centre | OF entries centre | OF latency | LDB distance | LDB velocity | LDB time light | LDB entries | LDB latency |
| --- | --- | --- | --- | --- | --- | --- | --- | --- | --- | --- |
| **OF distance** |  | 1 | 0.27 | 0.83 | -0.17 | 0.55 | 0.46 | 0.3 | 0.23 | 0.07 |
| OF velocity |  |  | 0.27 | 0.83 | -0.17 | 0.55 | 0.46 | 0.3 | 0.23 | 0.07 |
| **OF time centre** |  |  |  | 0.61 | -0.45 | 0.19 | 0 | 0.23 | 0.11 | -0.04 |
| OF entries centre |  |  |  |  | -0.27 | 0.51 | 0.38 | 0.31 | 0.2 | 0.05 |
| **OF latency** |  |  |  |  |  | 0.01 | 0.09 | -0.08 | 0 | 0.11 |
| LDB distance |  |  |  |  |  |  | 0.6 | 0.71 | 0.51 | -0.2 |
| LDB velocity |  |  |  |  |  |  |  | -0.12 | 0.38 | -0.01 |
| **LDB time light** |  |  |  |  |  |  |  |  | 0.28 | -0.2 |
| **LDB entries** |  |  |  |  |  |  |  |  |  | -0.41 |
| **LDB latency** |  |  |  |  |  |  |  |  |  |  |

| FEMALES | OF distance | OF velocity | OF time centre | OF entries centre | OF latency | LDB distance | LDB velocity | LDB time light | LDB entries | LDB latency |
| --- | --- | --- | --- | --- | --- | --- | --- | --- | --- | --- |
| **OF distance** |  | 1 | 0.46 | 0.89 | -0.17 | 0.56 | 0.48 | 0.25 | 0.56 | -0.27 |
| OF velocity |  |  | 0.46 | 0.89 | -0.17 | 0.56 | 0.48 | 0.25 | 0.56 | -0.27 |
| **OF time centre** |  |  |  | 0.71 | -0.2 | 0.33 | 0.19 | 0.25 | 0.41 | -0.24 |
| OF entries centre |  |  |  |  | -0.22 | 0.48 | 0.41 | 0.22 | 0.48 | -0.29 |
| **OF latency** |  |  |  |  |  | -0.1 | -0.02 | -0.13 | 0.01 | 0.11 |
| LDB distance |  |  |  |  |  |  | 0.61 | 0.66 | 0.65 | -0.41 |
| LDB velocity |  |  |  |  |  |  |  | -0.18 | 0.4 | -0.13 |
| **LDB time light** |  |  |  |  |  |  |  |  | 0.41 | -0.38 |
| **LDB entries** |  |  |  |  |  |  |  |  |  | -0.49 |
| **LDB latency** |  |  |  |  |  |  |  |  |  |  |

